# Oceanographic drivers of legal-sized male Dungeness crab in the California Current System

**DOI:** 10.1101/2023.03.05.531178

**Authors:** Ridouan Bani, André E. Punt, Daniel S. Holland, Nick Tolimieri, Kate Richerson, Melissa A. Haltuch, Nathan J. Mantua, Kiva L. Oken

## Abstract

We investigate environmental drivers of pre-season abundance of US West Coast legal-sized male Dungeness crab (Cancer magister), with the goal of developing an environmental index that can be used to forecast crab abundance in advance of the fishery. A conceptual life history approach is used to generate life-stage-specific and spatio-temporally-specific mechanistic hypotheses regarding oceanographic variables that influence survival at each life stage. Linear models are fit using the logarithms of pre-season abundance estimates of the coastal population of legal-sized male Dungeness crab as the dependent variable and environmental drivers from outputs developed using a regional oceanographic model for the California Current System as the independent environmental variables. Using different model selection methods, we show that the so-called ‘best’ models differ substantially among model selection approaches, illustrating the need to carefully choose performance metrics for model selection. Since our goal was to forecast crab abundance, we selected the ‘best’ model using a cross-validation metric that accounts for the time-series nature of the data. The resulting ‘best’ models suggest that the mechanisms that drive preseason abundance differ among regions widely recognized for spatially and seasonally varying dominant physical processes. We found that the processes determining pre-season abundance of legal-sized male Dungeness crab could be identified with sufficient precision to enable a predictive skill, suggesting that the predictions may be useful for management purposes. Moreover, we found that transport (within and between regions), as well as temperature were likely drivers of pre-season abundance, highlighting that future studies should focus on multiple processes.

## Introduction

The fishery for Dungeness crab (*Cancer magister* or *Metacarcinus magister*) is one of the most economically important on the US West Coast, with the value of Dungeness crab landings from the states of Washington, Oregon, and California combined averaging over $200 million annually in recent years (Pacific States Marine Fisheries Commission, 2019). During the 2019 season, Dungeness crab represented approximately 40% of the total ex-vessel revenue off the US West Coast, while representing only about 9% of the total catch biomass. In addition, there are high levels of cross-participation between the Dungeness fishery and other fisheries, such that changes in crab availability and regulations impacts participation and revenues in other fisheries (Fuller et al., 2017; Holland and Leonard, 2020; Fisher et al., 2021).

The Dungeness crab populations off the US West Coast are a single evolutionary population (Jackson et al., 2018) distributed throughout the California Current Marine Ecosystem and spanning nearly 3,000 km of coast from Southern British Columbia in Canada to Baja, Mexico. US West Coast habitats constitute approximately the southern third of the Dungeness crab range that extends north to the Pribilof Islands, Alaska. The Dungeness crab fishery is managed by the US West Coast states under the 3S management scheme: size, sex, and season. Males with a carapace widths >= 159 mm may be harvested during the open season, which generally runs from mid-winter through summer. The three US West Coast states (Washington, Oregon, and California) consult under the Pacific States Marine Fisheries Commission (PSMFC) Dungeness Crab Tri-state Process on issues affecting the commercial Dungeness crab fishery such as procedures for pre-season meat recovery testing and opening and closing days.

The trends of Dungeness crab-vary among regions, states, and years. The large majority of legal-sized males are harvested each year, and estimated abundance of pre-season legal-sized males follows the trends of the total catch, indicating that catch is a good proxy for abundance in most years (Richerson et al., 2020). Before 1980, the abundance of legal-sized male Dungeness crab in Northern California (north of 38.77°), Oregon, and Washington followed decadal cycles, and abundance in Central California (south of 38.77°) fluctuated independently with weak periodicity (Botsford et al., 1982). After 1980, the trends in legal-sized male Dungeness crab abundance changed and do not appear to exhibit strong periodicity (Rasmusson, 2013; Richerson et al., 2020).

Understanding the causes and consequences of fluctuations in Dungeness crab abundance and forecasting recruitment is important for fishers, processors, and managers of Dungeness crab and other fisheries to which crab fishers divert effort as crab catch rates decline during each season (Oken et al., 2020; Holland and Leonard, 2020; Fisher et al., 2021). In general, marine populations fluctuate due to 1) varying abiotic environmental conditions (Moran, 1953), 2) biotic interactions with other species (e.g., prey and predators), 3) fishing (Shelton et al., 2011), 4) dispersal (Bani et al., 2019), 5) nonlinear dynamics (Munch et al., 2020), and 6) other eco-evolutionary processes (Waples et al., 2010). The causes of fluctuations in Dungeness crab populations are not fully understood and continue to be the subject of ongoing research, although some studies point to environmental conditions during the dispersal phase and the nonlinearity in dynamics due to cannibalism as two main causes of fluctuations (Rasmuson, 2013; Shanks, 2013; Higgins et al., 1997; Botsford, 1986). There is no apparent stock-recruit relationship in this species (Shanks and Roegner, 2007), so the abundance of spawning adults is unlikely to directly impact recruitment.

Initial research on the fluctuations in Dungeness crab abundance focused on the effects of environmental conditions on dispersal by correlating lagged commercial catch to physical factors such as upwelling (Botsford and Wickham, 1975; Peterson, 1973), wind (Johnson et al., 1986), water temperature (Wild, 1980, Botsford and Lawrence, 2002), and transport (McConnaughey et al., 1992, Hobbs et al., 1992). The number of megalopae, the last larval stage of Dungeness crab, caught by a light trap placed in a marina in Charleston, Oregon is correlated with the spring transition (Shanks and Roegner, 2007) and internal tides (Rasmuson et al., 2020). Furthermore, the number of megalopae is correlated with the commercial catch in Oregon four years later (Shanks, 2013). The correlation between environmental factors and the commercial catch of Dungeness crab (Botsford and Wickham, 1975; Peterson, 1973; Johnson et al., 1986; Wild, 1980; Botsford and Lawrence, 2002; McConnaughey et al., 1992; Hobbs et al., 1992) or megalopae (Shanks and Reogner 2013; Rasmusson et al., 2020; Shanks 2013) allows for some degree of prediction of the Oregon commercial catch four years ahead (Shanks et al., 2010; Shanks, 2018). California and Washington catches are not closely correlated with those from Oregon, and predictions have not been attempted. However, it is possible that the effects of environmental factors on recruitment that appear to manifest at later stages such as the megalopae or legal-sized (i.e., commercial catch) stages could actually be due to effects at previous stages.

The shortest Dungeness crab life stage (a few seconds to minutes) is the pre-zoea (Buchanan and Milleman, 1969), which occurs in shallow nearshore waters (Stone and O’Clair, 2002; Scheding et al., 1999; Armstrong et al., 1988). After the pre-zoea stage, the zoea 1, 2, 3, 4, 5, and megalopae stages last about one to four weeks for each stage. As the larvae age, their spatial extent increases such that stage 5 zoea can be found at greater distances latitudinally and offshore than earlier stages (Reilly, 1983a). The varying spatial and temporal distributions mean that different environmental factors likely impact survival at each life stage. Also, crab larvae originating from different locations along the coast are subject to varying conditions that can drive substantial spatial differences in recruitment spatially and temporally.

Environmental relationships have been incorporated into stock assessments to quantify recruitment variability that can be attributed to environmental covariates and to evaluate the impact of projected climate change on stocks (Maunder and Thorson, 2019; Haltuch and Punt, 2011; Schirripa et al., 2009). It has been suggested that environmental indexes need to be able to explain at least 50 % of the total variation in recruitment to be meaningful for management (de Oliveira and Butterworth, 2005). To achieve this, it is necessary to develop frameworks to match the temporal and spatial distributions of multiple life-stages and environmental and biological forcing. This is especially the case during the larval phase where there are many pre-recruit life-stages of short duration, each with different levels and sources of vulnerability. However, empirical statistical relationships between environmental drivers and population dynamics may not hold up over time. Informing statistical models with a mechanistic understanding of physical-biological relationships may improve predictive capacity (Jacox et al., 2020)

Recently, coupled biophysical modeling methods have been used to investigate the exposure history of individual Dungeness crab and other marine species during the pelagic and adult phases (Norton et al., 2020; Haltuch et al., 2020; Tolimieri et al., 2018). These new methods, although computationally expensive, are more realistic and offer new insights regarding the interactions among biological, chemical, and environmental factors and species life-cycle stages. For example, Norton et al. (2020) used oceanographic factors extracted from the Joint Institute for the Study of the Atmosphere and Ocean (JISAO) Seasonal Coastal Ocean Prediction of Ecosystem (J-SCOPE) model, a high-resolution coupled physical-biogeochemical model, to identify the exposure history that affects the number of megalopae collected during surveys conducted by NOAAs Northwest Fisheries Science Center (NOAA/NWFSC) at 37 stations off the Washington and Oregon coasts from 2009 to 2017. However, Norton et al. (2020) only investigated the exposure history of Dungeness crab during the megalopae stage, which is the last larval stage, whereas we consider all the life-cycle stages and for California as well as Washington and Oregon.

While Dungeness crab larvae develop between January and July, the ocean currents along the West Coast of US undergo seasonal and decadal variation at various spatial scales (Checkley and Barth, 2009). Water temperature is the only factor known to affect Dungeness crab throughout the female conditioning, spawning, pelagic, and juvenile phases (Wild 1980; Sulkin and McKeen 1989; Rasmussen 2013; Norton et al., 2020). Bottom temperature can affect metabolism, allocation of energy, reproductive output, quality of eggs, the chances of successful spawning, as well as survival of juveniles. The temperature at the top of the water column can affect the development and survival of the larvae. Other environmental factors such as salinity, pH, and hypoxia can impact the survival rate, but to a lesser extent than temperature (Rasmussen 2013). Adult Dungeness Crab are likely less sensitive to environmental conditions than younger life stages, and they can move to avoid areas of lower salinity and hypoxic conditions (Bernatis et al., 2007; Curtis and McGae 2008; Froehlich et al., 2014).

We developed a conceptual life history approach to identify life-stage-specific and spatio-temporally specific mechanistic hypotheses regarding oceanographic variables that influence survival at each life stage of Dungeness crab. We then modeled the estimated pre-season abundance of the coastal population of legal-sized male Dungeness crab from 1970 to 2016 (Northern and Central California) and from 1982-2016 (Oregon and Washington) (Richerson et al., 2020) as a function of environmental drivers from Regional Ocean Modeling System (ROMS) outputs for the California Current System using General Linear Models (GLMs). ROMS is a free-surface, terrain-following, primitive equations ocean model widely used for a diverse range of applications (Di Lorenzo, 2003; Wilkin et al., 2005). The framework we use is an expansion of previous methods applied to sablefish *(Anoplopoma fimbria)* (Tolimieri et al., 2018) and petrale sole *(Eopsetta Jordani)* (Haltuch et al., 2020).

## Methods

### Overview and data

The dependent variables are time-series of log-transformed estimates of pre-season abundance of legal-sized male Dungeness crab (Central and Northern California, 1970-2016, and Oregon and Washington, 1982-2016) that are analyzed regionally because the Dungeness crab fisheries are managed by state, and because there are regional differences in (i) the date the fishing season opens, (ii) biological traits such as growth rate, spawning, and larval duration, and (iii) trends in pre-season abundance (Fig. 1). Richerson et al. (2020) estimated pre-season abundance of legal-sized male Dungeness crab and catchability by applying a Bayesian version of the traditional linear depletion estimator (Leslie and Davis, 1932) to data from fish tickets (i.e., landing receipts) and fisher’s logbooks. California was divided into Northern and Central California regions, with the division at the Sonoma/Mendocino County line (38.77°N), consistent with fishery management practices. There is little Dungeness crab fishing activity in Southern California, so this region is omitted

**Figure 1:**
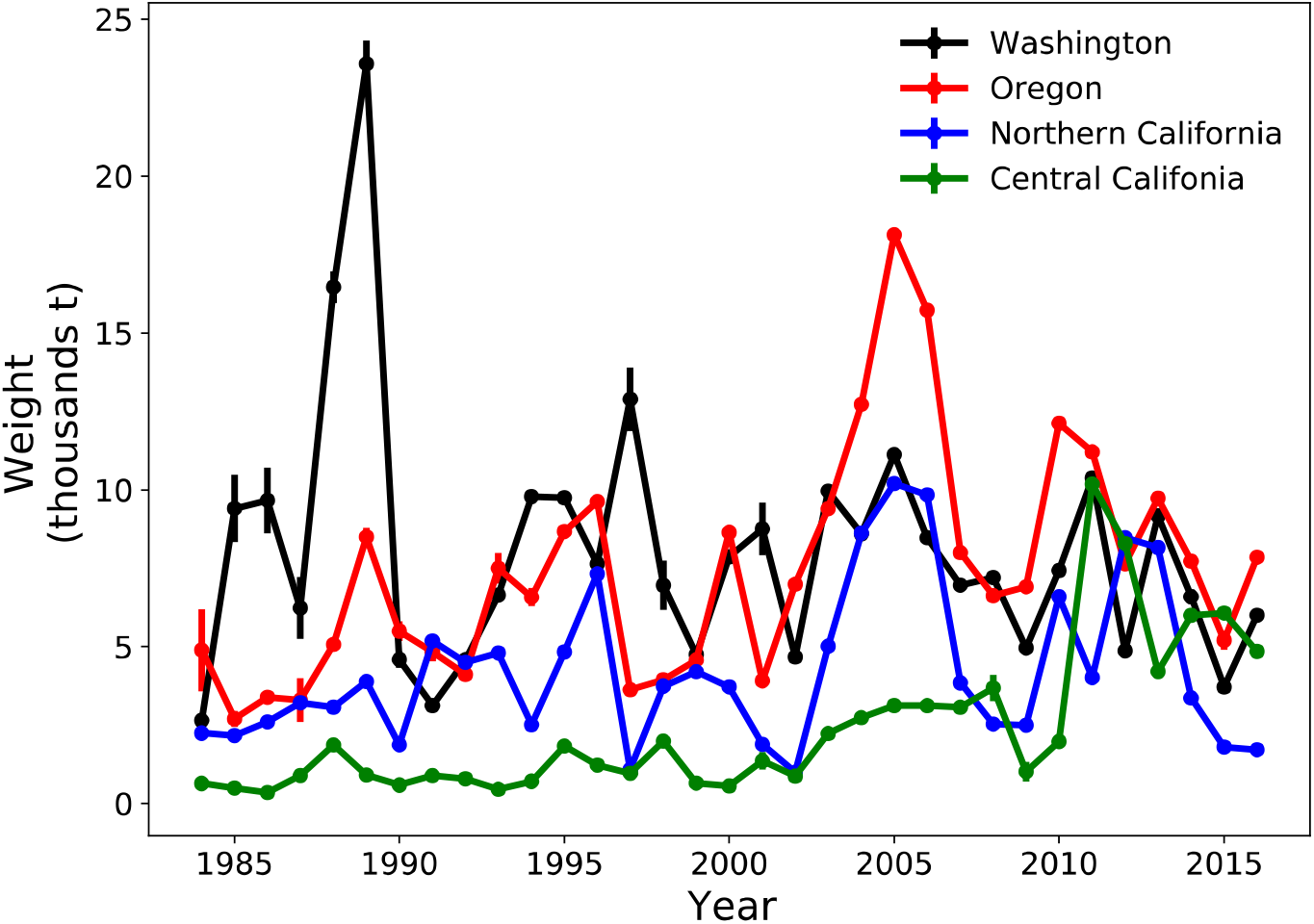
Preseason abundance of legal-sized male Dungeness crab by region and year. The lines and bars indicate mean and standard deviation of estimated preseason abundance (Richerson et al., 2020).

For each region, we identified life-stage-specific and spatiotemporally specific mechanistic hypotheses regarding oceanographic variables (i.e., independent variables) that may influence Dungeness crab survival and recruitment to suitable habitats at each life stage. The oceanographic variables were obtained from a California Current Ecosystem configuration of the ROMS with 4-Dimensional Variational (4D-Var) data assimilation (Neveu et al., 2016) (https://oceanmodeling.ucsc.edu). The ROMS domain covered the region 30–48°N and from the coast to 134°W at 0.1° (~10 km) horizontal resolution, with 42 terrain-following vertical levels. The ROMS’s 1980-2010 historical reanalysis uses 8-day overlapping analysis cycles where the start day corresponds to day 4 of the previous cycle. New initial conditions are used during each cycle.

All oceanographic independent variables were 4-day averages, summarized as either the average or the maximum over the spatial and depth extents selected for each life stage (Fig. S1). Near-surface and bottom layers follow the ROMS’ 42 vertical layers. Larval stages tend to be found in near-surface waters (Jamieson and Phillips, 1988), which can be approximated using the ROMS’ top layer, and the adult and juvenile stages can be found at the bottom, which can be approximated using the ROMS’ bottom layer. The widths of the vertical layers in ROMS increase with sea depth, which means that the top layer in ROMS is within the mixed layer especially within the continental shelf (depth 0-500 m) and up to depths greater than 2,000 m (2,000m/41layer=48.78m/layer), assuming that the surface mixed layer is between the sea surface and 50m depth.

We examined the choice of the functional forms between the dependent and independent variables by exploring the use of quadratic terms. We also explored how different model selection approaches influence the final models and their performance using several performance metrics.

### Identification of hypotheses

We reviewed the available literature on the life history of Dungeness crab (see Rasmussen (2013) for a full review) and identified the ecological, biological, and environmental processes that can affect growth, survival, and successful recruitment to suitable habitats. Successful recruitment of 4-year-old males into the fishery is the result of a series of events from female preconditioning and spawning through the four years needed to grow to harvestable size and can be impacted by several processes. We identified hypotheses based on environmental conditions, match/mismatch with food, prey and predator interactions, and intra-specific competition including cannibalism. The initial set of hypotheses had to be restricted owing to lack of data to quantify some of them. Consequently, the 27 hypotheses considered in the quantitative analyses were largely related to the effects of oceanographic conditions on survival and physical transport of Dungeness crab, consistent with previous research (Table 1).

**Table 1:**
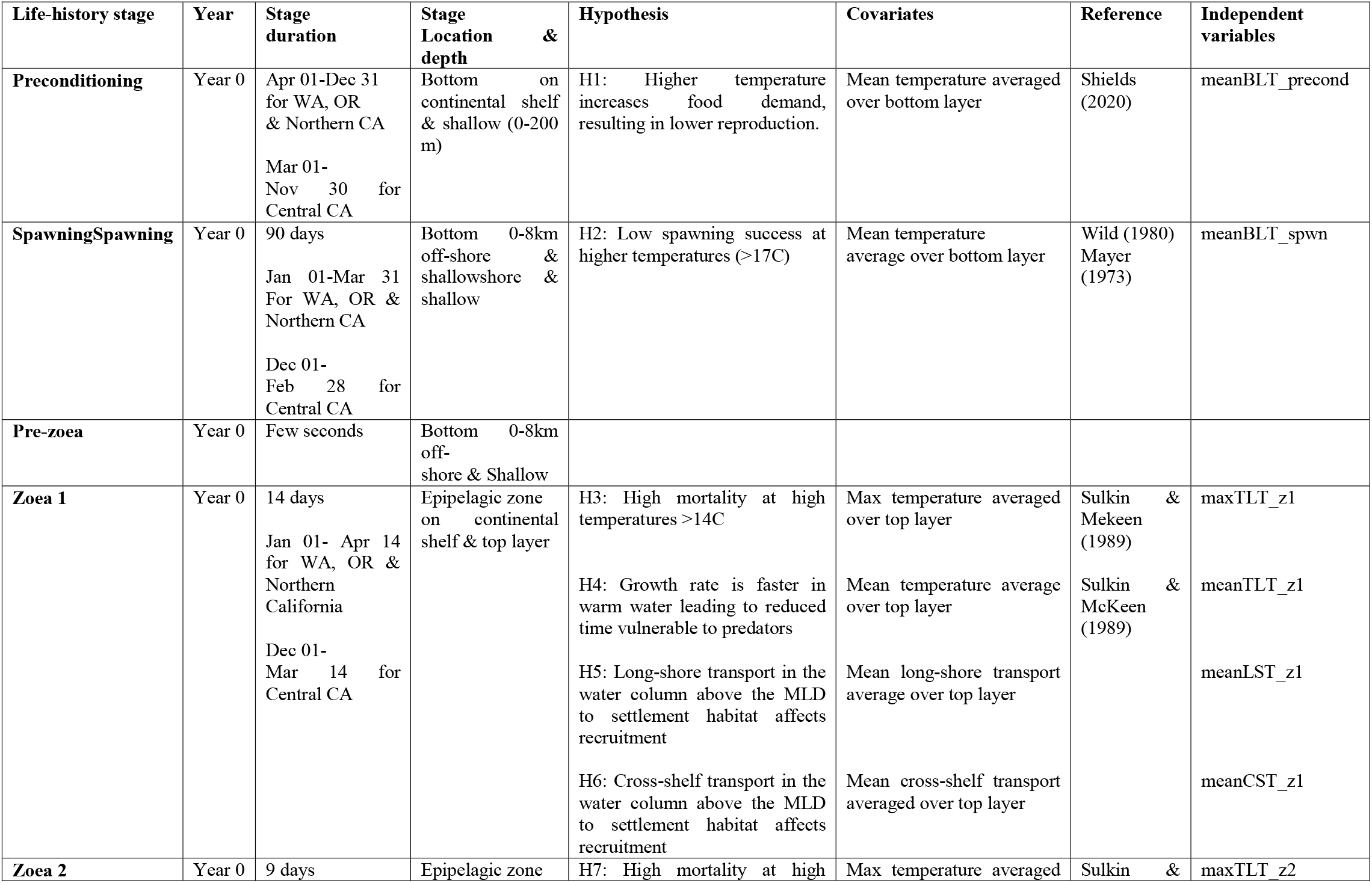
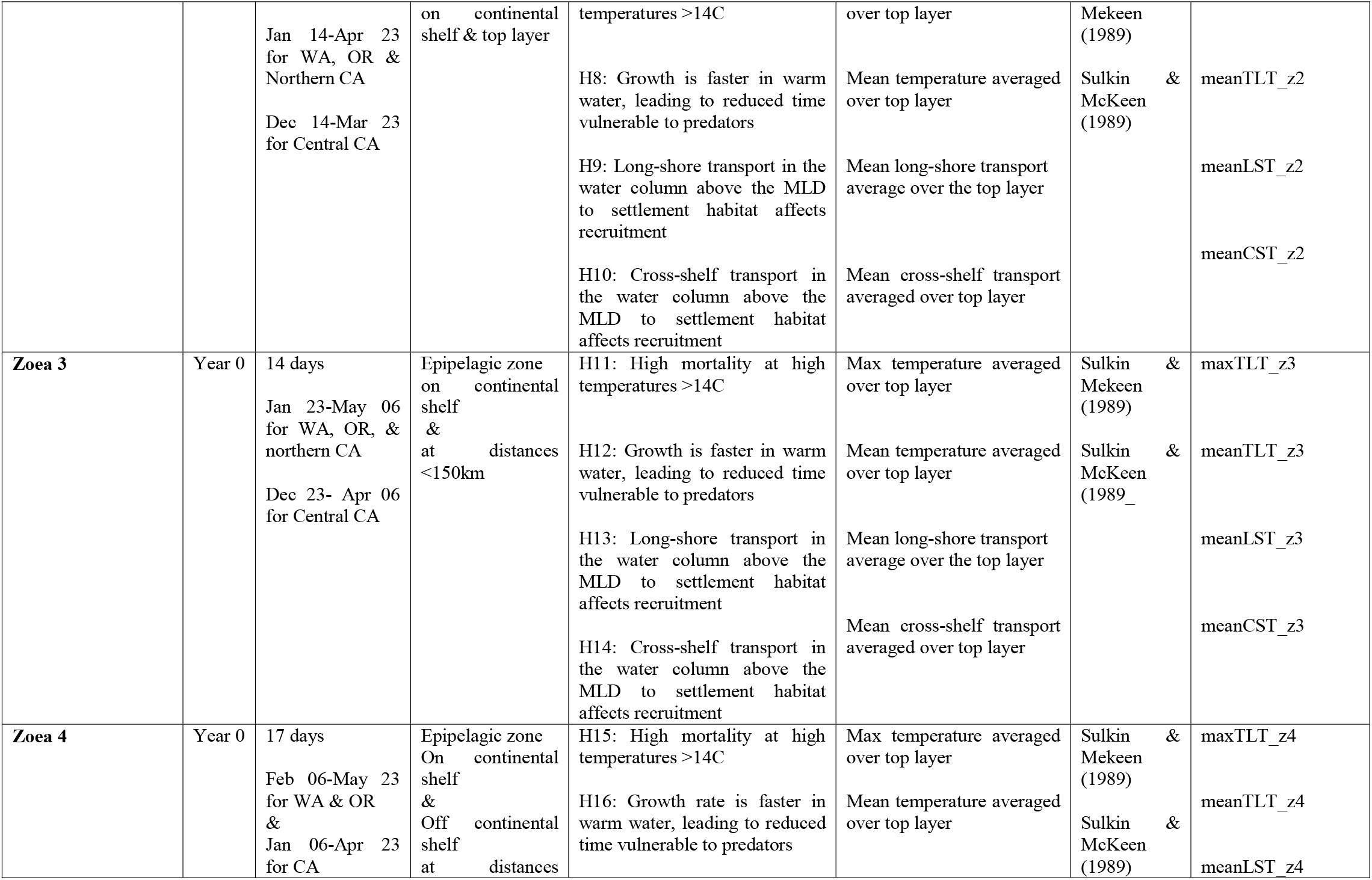
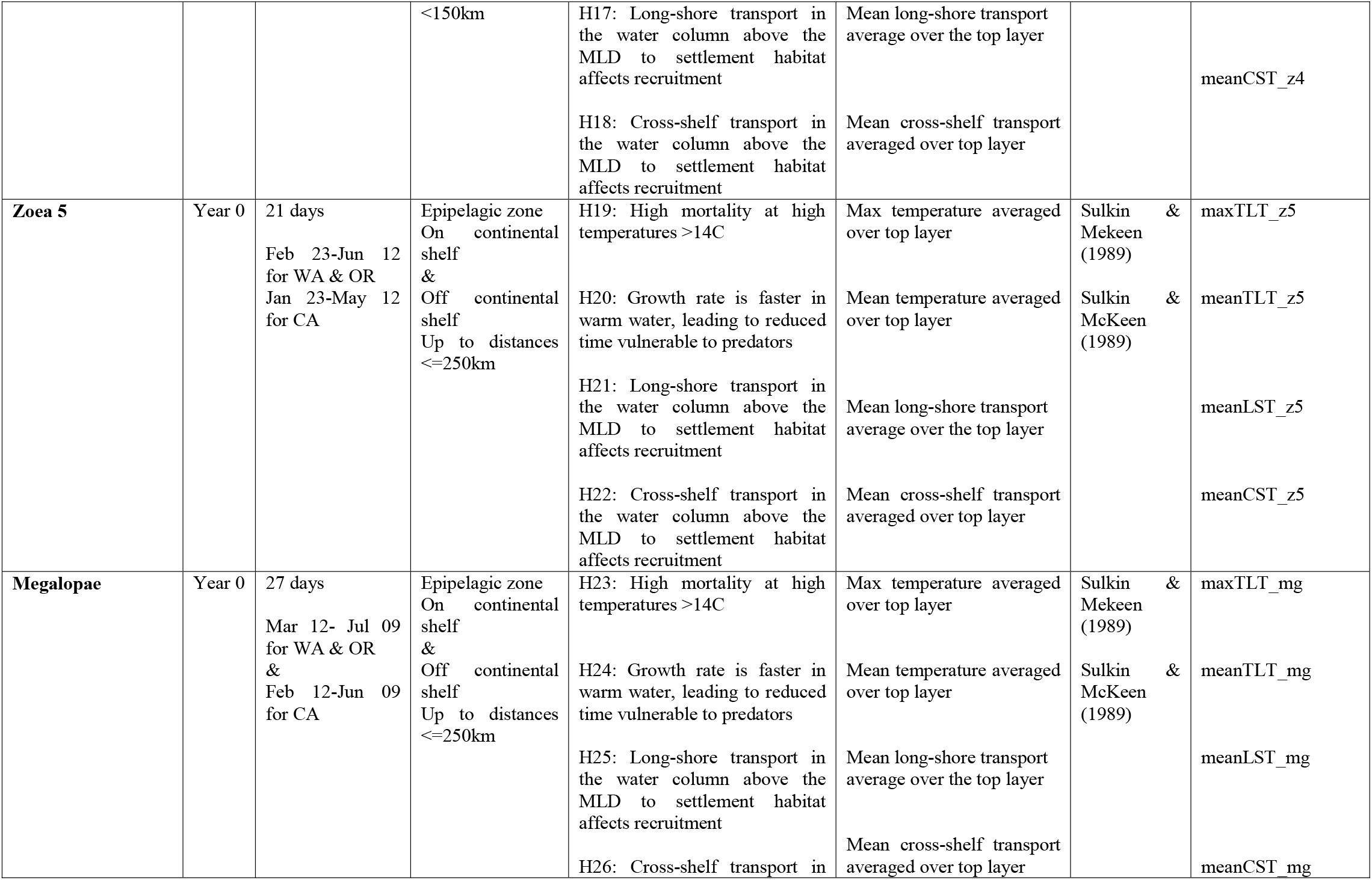
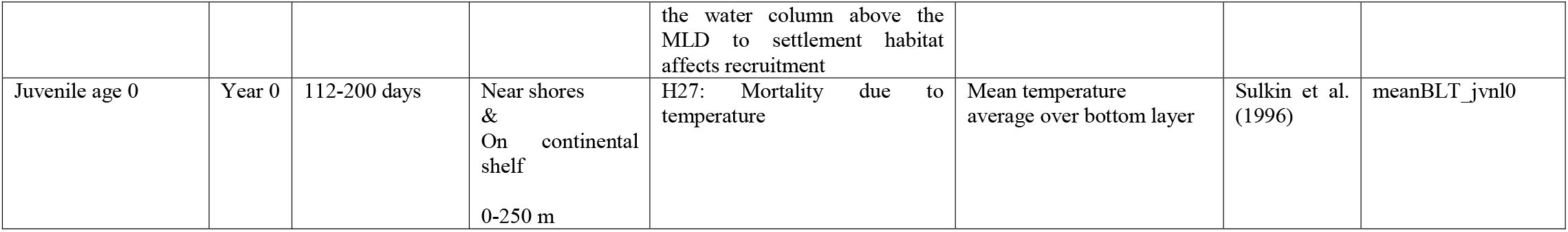
Conceptual life history models by region (Washington, Oregon, and Northern and Central CA)

### Modeling

The modelling was conducted separately for each of the four regions. Model selection was based on cross validation, corrected Akaike Information Criterion (AICc), and explained variance, followed by a series of validation tests to evaluate whether data points most impacted the selected best-models and their predictive skill. We ran individual linear and quadratic regressions for each independent variable against the natural logarithms of Dungeness crab pre-season abundance to examine whether the data supported non-linear over linear relationships. Regressions were based on the independent variables from ROMS as well as alternative independent variables computed by conducting a principal component analysis on the original independent variables and then selecting the five principal components that explained the most variation in the original independent variables as independent variables. We evaluated the correlations between independent variables and excluded models with high multicollinearity. Finally, we tested for autocorrelations in our best-fit models and corrected for them when significant.

### Model selection

Generalized linear models (GLMs) were run for all combinations of up to five independent variables (27! / ((27-5)!5!) =80,730 possible models). The restriction to five independent variables (i.e., one for every six data points in the time series) was to avoid overfitting. All possible models were tested for collinearity using the Pearson correlation coefficient. Models with at least one absolute value of pairwise correlation greater than 0.75 were eliminated from the model selection process. After checking for collinearity, 27,530, 13,731, 28,815, and 5,934 models remained for Washington, Oregon, Northern California, and Central California, respectively. The model selection involved evaluating and ranking and then comparing the performances of the remaining models using the following five metrics:

1. ΔAICc, the difference between each model and that with lowest AICc on the whole data set.
2. Calculating the 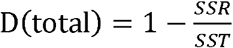, where the sum of squares total, *SST*, is the squared differences between the observed dependent variable and its mean, and the sum of squared residuals, *SSR,* is the sum of squared differences between the observed dependent variables and their predicted values using the model fitted to the whole time series (Fig. 2a).
3. Creating 1,000 pseudo data sets by randomly dividing the original data into training (80%) and testing (20%) data sets and fitting each model to the training data set and then calculating the average 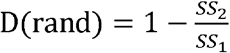, where *SS*_1_ is the sum of squared differences between the observed dependent variables from the testing data set and its mean from the training set, and *SS_2_* is the sum of squared differences between the observed dependent variables from the testing set and their predicted values using the model fitted to training data (Fig. 2b,c).
4. Fitting all the models to seven temporally ordered sampled sets (blocks) representing 80% of the data and then calculating the average 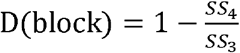, where *SS*_3_ is the sum of squared differences between the observed dependent variables from the predicted 20% of the data set and its mean from the 80% of the data used for model fitting, and *SS*_4_ is the sum of squared differences between the observed dependent variables from the predicted 20% of the data and their predicted values using the model fitted to the 80% of the data (Fig. 2d,e).
5. Fitting all the models to the data for 1985-2009 and then calculating 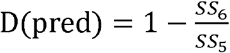, where *SS*_5_ is the sum of squared differences between the observed dependent variables from 2010-2014 and its mean from 1985-2009, and *SS_6_* is the sum of squared differences between the observed dependent variables from 2010-2014 data and their predicted values using the model fitted to 1985-2009 data (Fig. 2f).

**Figure 2:**
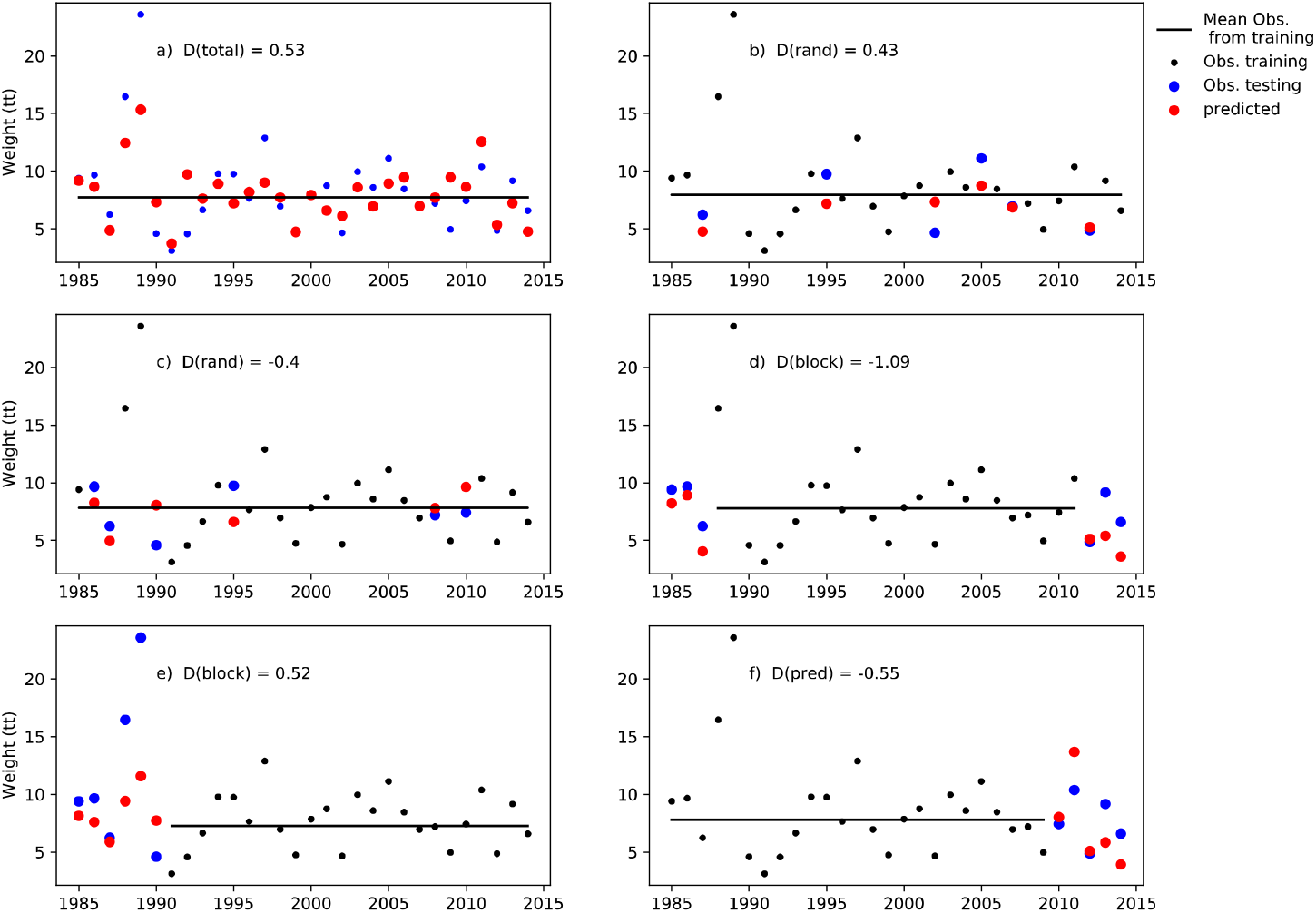
Using the Washington preseason abundance of legal-male Dungeness crab, we illustrate the methods used to calculate D(total) in plot a), D(rand) in plots b) and c), D(block) in plots d) and e), and D(pred) in plot f). The training set (black dots), their average (black straight line), the testing set (blue dots), and testing set predicted values using the model fitted to training data (red dots). In a) the training set is also the testing set, in b) and c) we randomly sampled the testing set twice, in d) and e) the training set is two samples of ordered data, and in f) the testing set is the last five years. For example, in plot c) D(rand) = −0.4 and graph d) D(block) = −1.09; the sum of the squared differences between the red and blue dots is greater than the sum of the squared differences between the black dots and the black straight line, meaning that the average values of training sets can predict the testing sets better than predicted values using the model fitted to training sets. Compared to graph c) D(rand) = 0.78 and graph d) D(block) = 0.52 where the predicted values using the best model fitted to training data can predict the testing set better than the average value from the training set.

The metrics measure how well models explain the observed data (D(total), and ΔAICc), how models will perform given new data (D(rand)), how models will perform on new data given the potential autocorrelation among recruitments (D(block)), and how models will perform on new data in the future given potential autocorrelation (D(pred)). Note that the data on which D(pred) is based is among the seven blocks used to calculate D(block) since 2009-2014 is a sample of ordered points. Although these measures of predictive skill are similar in form to an r-square, except for D(total), they are not a measure of the variance explained in the holdout sample but rather the amount of variance explained relative to a prediction based on just using the mean from the training sample (e.g., how much better can we do than just assuming recruitment will equal the observed mean in previous years).

We repeated the model selection process including the one-year and two-year lagged response variables as independent variables to account for potential autocorrelation in the response variable, and examined whether models with lagged response variables perform better than those without lagged response variables. Generalized linear models (GLMs) were run for all combinations of up to five independent variables including the one-year and two-year lagged response variables (29! / ((29-5)!5!) =118,755 possible models). After checking for collinearity, 50,719, 26,983, 49,170, and 11,693 models remained for Washington, Oregon, Northern, and Central California, respectively. The model selection involved evaluating and ranking the models and then comparing the performances of the remaining models using ΔAICc, D(block), D(pred), and D(total).

### ‘Best’ model testing and diagnostics

The following analyses were conducted in addition to assessing the distribution of residuals, and homoscedasticity for the ‘best’ model.

1. A jackknife was conducted to assess the extent to which model predictions change if one year’s data are ignored and hence evaluate the extent to which the ‘best’ model was sensitive to influential years.
2. The ‘best’ model was used to fit the data for 1985-2008 and then predict the data for 2009-2014 to test ‘best’ model predictive skill.

## Results

### Life history and selection of independent variables

Female preconditioning covers the period between two successive spawning events (April-December in Washington, Oregon, and Northern California, and March-November in Central California). During the female preconditioning phase, females are found on shell habitats, specifically oyster beds (Dumbauld, 1993; Cleaver, 1949). Several months later, molted males and females are found in shallow waters nearshore (Rasmussen 2013). Higher water temperature can affect female preconditioning (Table 1, hypothesis 1, hereafter, H1), with females allocating less energy to reproduction when temperatures are higher, resulting in fewer eggs and/or a lesser egg quality (Shields 2020).

Females extrude their eggs, inseminate them, and then attach them to setae below the abdominal flap a few months following copulation (October-December in Washington, Oregon, and Northern California, and September-November in Central California). A female can carry between 1.5 and 2.5 million eggs (Hankin et al., 1989) and eggs are released when the pre-zoea are fully developed (December-February in Central California and January-March in Washington, Oregon, and Northern California (Rasmussen, 2013). Fewer eggs are produced, and hatching success declines when temperature rises from 13 to 17°C (Mayer, 1973) (Table 1, H2).

The survival and development of Dungeness crab larvae from the Puget Sound decreased with an increase in water temperature (Sulkin and McKeen, 1989). In addition, Sulkin and Mekeen (1989) found that stage-specific survival decreased when larvae were exposed to increasing temperature (Table 1, H3, H7, H11, H15, H19, and H23; zoea 1, 2, 3, 4, 5 and megalopae, respectively). Moreover, larval vulnerability to temperature increased with age. Water temperature can also affect survival of Dungeness crab larvae by increasing development speed and decreasing the time spent at each stage (Sulkin and McKeen, 1989). With sufficient food, the rapid development in warmer water can help the larvae outgrow potential predators (Table 1, H4, H8, H12, H16, H20, and H24).

Ocean currents transport Dungeness crab larvae, which in turn affect survival and settlement. Traditionally, studies relied on indexes such as upwelling, downwelling, wind direction, and the Pacific Decadal Oscillation (PDO) as indicators of physical transport. For example, Shanks (2018) argued that megalopae catches tend to be low when the PDO is positive and high when it is negative, potentially due to enhanced southward transport. We used two proxies for physical transport during the larval stage: the average: a) cross-shelf and b) long-shore transport of the top water column. To calculate these variables, we first calculated the projections of ROMS’ U- and V-momentum on the line between the center of each ROMS’ grid cell and closest point on the coast and on the line perpendicular to the line between the center of each ROMS’ grid cell and closest point on the coast. Then, the projections of U- and V-momentum on the line between the center of each ROMS’ grid cell and closest point on the coast were added to obtain the cross-shelf transport vector. The long-shore transport vector was obtained analogously.

We calculated the temporal and spatial averages of the cross-shelf transport and long-shore transport components (in meters per second) over the spatial extent by stage (Fig. S1). Positive cross-shelf (long-shore) transport values indicate currents are pushing larvae toward the shore (northward).

Zoea 1 larvae are released within 8-10 km offshore and migrate off the continental shelf as they develop (Reilly, 1983b) (Fig. S1). Zoea 1 and 2 are commonly found on the continental shelf (Table 1, H3-H10), and zoea 3 and 4 are found off the continental shelf at distances > 150 km (Reilly, 1983b). We assumed that zoea 3 and 4 occur on and off the continental shelf up to 150 km (Table 1, H11-H18) and zoea 5 and megalopae up to 250 km (Table 1, H19-H26). Megalopae are known to be stronger swimmers compared to other stages (Fernandez et al., 1994), and can be found to depths of 25m in the open ocean during daylight (Jamieson and Phillips, 1993).

### Selection of covariates

Correlations among independent variables varied widely and tend to be higher between variables with higher spatial or/and temporal overlap (Figs S2-S5, Tables S1-S4). Specifically, the correlations between the variables for zoeal stages 1 and 2, between zoeal stages 3 and 4, and between zoeal stages 5 and megalopae tended to be high, except for Northern California where the correlations between variables for zoeal stages 3 and 4 are low (Table S3).

Models were fit with each independent variable treated as linear and quadratic coefficients (Figs S6-S9). However, we did not find a significant difference in fit between the linear and quadratic forms. In some cases, a single outlier point can cause the R^2^ value of the quadratic form to be higher than the R^2^ of the linear form (e.g., the year 1993 in Central California, with a low maximum temperature corresponding to a high preseason abundance 4 years later, Fig. S9)).

The best-fitting models with five or fewer of the top five principal components did not perform as well as the original independent variables (Tables S5-S13) so the remaining analyses use only the original independent variables.

## Model selection

### Model’s performance depended on the data and region used to test the models

We hypothesized that the best models would be ranked similarly using ΔAICc, D(rand), D(block), D(pred) and D(total), but this was not the case (Tables 2–5). Only the Central California models selected using ΔAICc were the same as those selected using D(rand), and D(block), and the same models performed relatively well when tested against recent 2009-2014 data (D(pred) = 0.86) and the entire (1985-2014) data set (D(total)= 0.71) (Table 5). The top ranked model in Central California fitted the testing data better than the historical training data’s average when fitted to any randomly sampled test data set (D(rand) = 0.55) and contiguous blocks of five years of data (D(block) = 0.73) (Table 5). The best models for Central California showed a larger improvement over the historical average of training data for predicting testing data compared to Washington, Oregon, and Northern California.

**Table 2:**
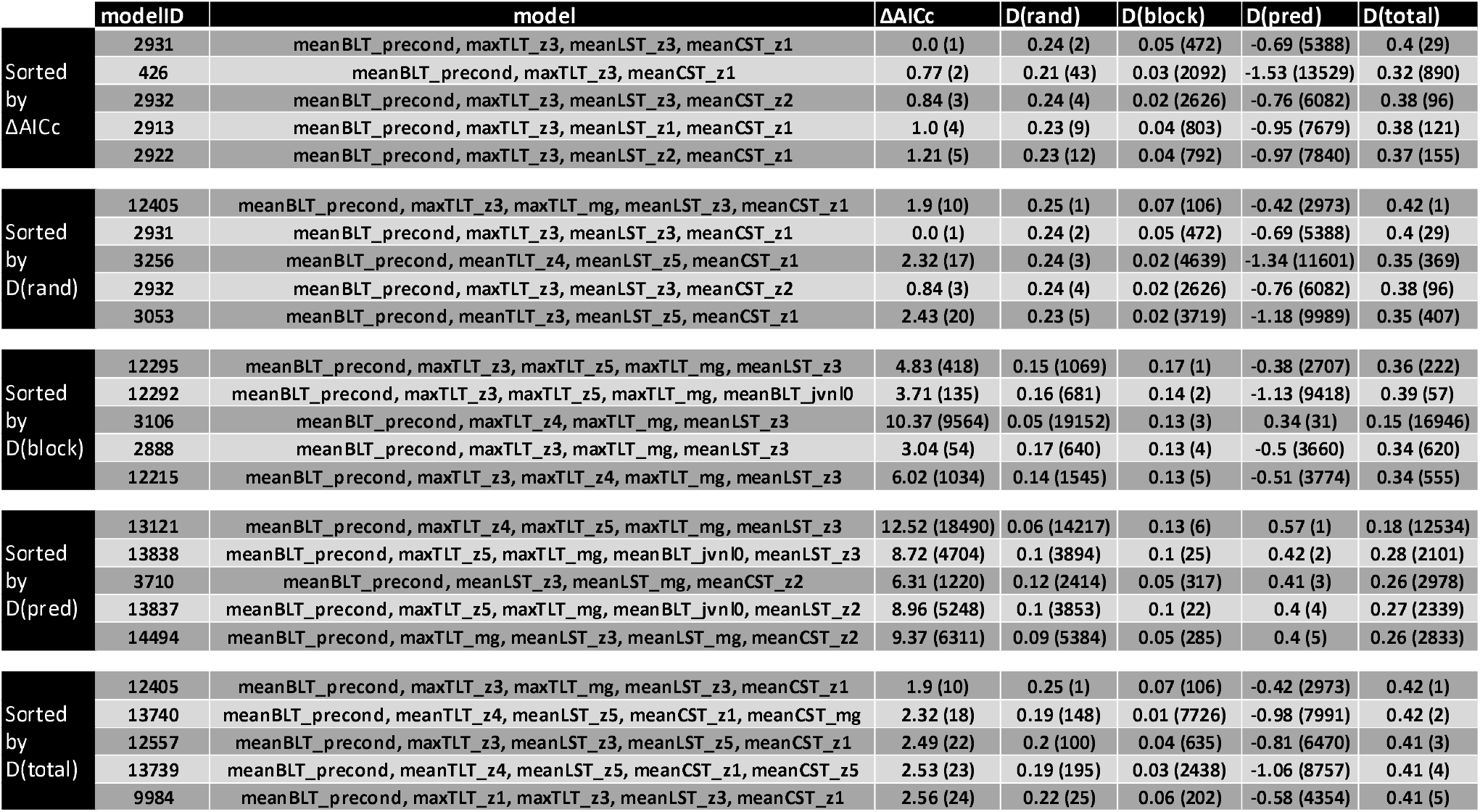
Top five models for each of the five metrics for Washington for the models without lagged variables. The results show the value of the metric and its rank (between parentheses). ΔAICc is ranked in increasing order and D(rand), D(pred), D(block) and D(total) are ranked in decreasing order.

**Table 3:**
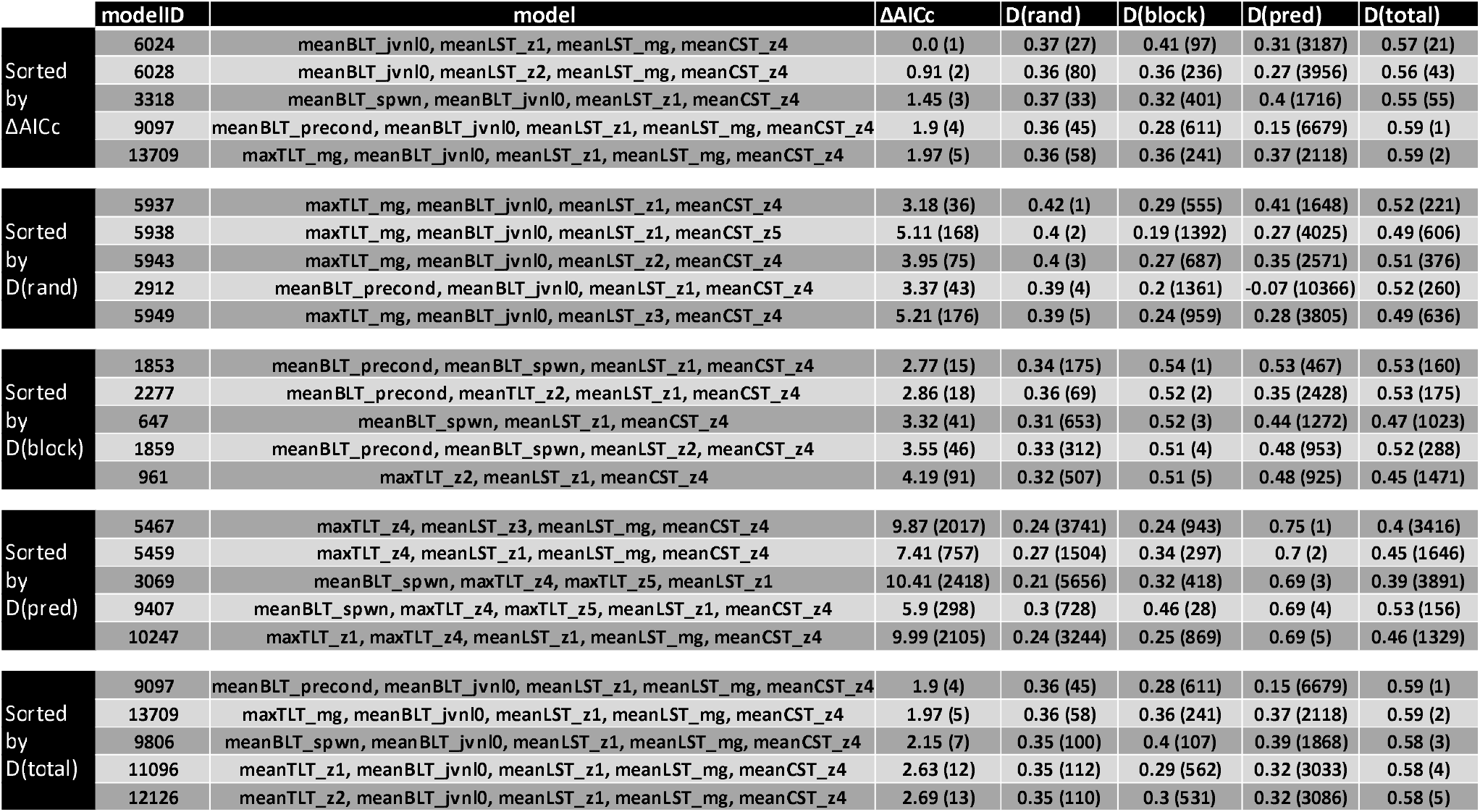
Top five models for each of the five metrics for Oregon for the models without lagged variables. The results show the value of the metric and its rank (between parentheses). ΔAICc is ranked in increasing order and D(rand), D(pred), D(block) and D(total) are ranked in decreasing order.

**Table 4:** Top five models for each of the five metrics for Northern California for the models without lagged variables. The results show the value of the metric and its rank (between parentheses). ΔAICc is ranked in increasing order and D(rand), D(pred), D(block) and D(total) are ranked in decreasing order.

| | modelID | model | $\Delta AICc$ | D(rand) | D(block) | D(pred) | D(total) |
| --- | --- | --- | --- | --- | --- | --- | --- |
| Sorted by $\Delta AICc$ | 1827 | meanTLT_z3, meanCST_z3, meanCST_z5 | 0.0 (1) | 0.41 (1) | 0.08 (14261) | 0.28 (19554) | 0.55 (233) |
|  | 8858 | meanTLT_z3, maxTLT_z5, meanCST_z3, meanCST_z5 | 0.61 (2) | 0.37 (11) | 0.04 (22000) | 0.15 (23120) | 0.59 (32) |
|  | 3311 | meanBLT_precond, meanTLT_z3, meanCST_z3, meanCST_z5 | 0.82 (3) | 0.4 (2) | 0.14 (9917) | 0.29 (18981) | 0.58 (38) |
|  | 182 | meanTLT_z3, maxTLT_z5 | 1.1 (4) | 0.35 (58) | 0.32 (1580) | 0.53 (6830) | 0.49 (3306) |
|  | 13899 | meanBLT_precond, meanTLT_z3, maxTLT_z5, meanCST_z3, meanCST_z5 | 1.18 (5) | 0.38 (5) | 0.1 (12065) | 0.17 (22672) | 0.62 (1) |
| Sorted by D(rand) | 1827 | meanTLT_z3, meanCST_z3, meanCST_z5 | 0.0 (1) | 0.41 (1) | 0.08 (14261) | 0.28 (19554) | 0.55 (233) |
|  | 3311 | meanBLT_precond, meanTLT_z3, meanCST_z3, meanCST_z5 | 0.82 (3) | 0.4 (2) | 0.14 (9917) | 0.29 (18981) | 0.58 (38) |
|  | 9109 | meanTLT_z3, meanLST_z4, meanCST_z3, meanCST_z5 | 1.25 (7) | 0.4 (3) | 0.11 (11510) | 0.29 (19130) | 0.58 (63) |
|  | 4405 | meanBLT_spwn, meanTLT_z3, meanCST_z3, meanCST_z5 | 2.53 (25) | 0.39 (4) | 0.07 (15329) | 0.3 (18369) | 0.56 (167) |
|  | 13899 | meanBLT_precond, meanTLT_z3, maxTLT_z5, meanCST_z3, meanCST_z5 | 1.18 (5) | 0.38 (5) | 0.1 (12065) | 0.17 (22672) | 0.62 (1) |
| Sorted by D(block) | 1602 | maxTLT_z3, meanTLT_z5, meanLST_z3 | 9.86 (6441) | 0.31 (464) | 0.5 (1) | 0.7 (285) | 0.38 (19375) |
|  | 5117 | maxTLT_z1, maxTLT_z2, meanTLT_z5, meanLST_z3 | 14.11 (18626) | 0.29 (1501) | 0.5 (2) | 0.71 (127) | 0.35 (21590) |
|  | 1290 | maxTLT_z2, meanTLT_z5, meanLST_z3 | 11.21 (10374) | 0.29 (1497) | 0.5 (3) | 0.71 (128) | 0.35 (21588) |
|  | 940 | maxTLT_z1, meanTLT_z5, meanLST_z3 | 11.21 (10373) | 0.29 (1500) | 0.5 (4) | 0.71 (129) | 0.35 (21589) |
|  | 7182 | maxTLT_z2, meanTLT_z5, maxTLT_mg, meanLST_z3 | 14.03 (18458) | 0.24 (7797) | 0.49 (5) | 0.7 (171) | 0.35 (21471) |
| Sorted by D(pred) | 1745 | meanTLT_z3, meanTLT_z5, meanCST_z3 | 3.78 (101) | 0.3 (810) | 0.34 (1027) | 0.75 (1) | 0.49 (3316) |
|  | 3064 | meanBLT_precond, maxTLT_z3, meanTLT_z4, meanCST_z3 | 10.24 (7474) | 0.26 (4852) | 0.15 (9083) | 0.75 (2) | 0.43 (14307) |
|  | 13346 | meanBLT_precond, maxTLT_z3, meanTLT_z4, meanLST_z4, meanCST_z3 | 13.33 (16661) | 0.23 (11795) | 0.13 (10274) | 0.75 (3) | 0.43 (14164) |
|  | 1717 | meanTLT_z3, meanTLT_z4, meanCST_z3 | 5.49 (426) | 0.27 (3516) | 0.36 (678) | 0.75 (4) | 0.46 (8103) |
|  | 3092 | meanBLT_precond, maxTLT_z3, meanTLT_z5, meanCST_z3 | 7.96 (2314) | 0.3 (588) | 0.21 (6133) | 0.74 (5) | 0.47 (6597) |
| Sorted by D(total) | 13899 | meanBLT_precond, meanTLT_z3, maxTLT_z5, meanCST_z3, meanCST_z5 | 1.18 (5) | 0.38 (5) | 0.1 (12065) | 0.17 (22672) | 0.62 (1) |
|  | 14084 | meanBLT_precond, meanTLT_z3, meanTLT_mg, meanCST_z3, meanCST_z5 | 1.48 (9) | 0.37 (15) | 0.16 (8483) | 0.29 (19141) | 0.62 (2) |
|  | 13965 | meanBLT_precond, meanTLT_z3, meanTLT_z5, meanCST_z3, meanCST_z5 | 1.64 (11) | 0.36 (22) | 0.17 (7815) | 0.32 (17760) | 0.61 (3) |
|  | 13900 | meanBLT_precond, meanTLT_z3, maxTLT_z5, meanCST_z3, meanCST_mg | 2.01 (15) | 0.38 (6) | 0.2 (6454) | 0.24 (20877) | 0.61 (4) |
|  | 28057 | meanTLT_z3, meanTLT_mg, meanLST_z4, meanCST_z3, meanCST_z5 | 2.43 (22) | 0.36 (23) | 0.1 (11994) | 0.32 (17744) | 0.6 (5) |

**Table 5:** Top five models for each of the five metrics for Central California for the models without lagged variables. The results show the value of the metric and its rank (between parentheses). ΔAICc is ranked in increasing order and D(rand), D(pred), D(block) and D(total) are ranked in decreasing order.

| | modelID | model | $\Delta AICc$ | D(rand) | D(block) | D(pred) | D(total) |
| --- | --- | --- | --- | --- | --- | --- | --- |
| Sorted by $\Delta AICc$ | 2074 | meanBLT_spwn, maxTLT_z1, maxTLT_z4, meanLST_mg | 0.0 (1) | 0.55 (1) | 0.73 (1) | 0.86 (7) | 0.71 (10) |
|  | 2077 | meanBLT_spwn, maxTLT_z1, maxTLT_z4, meanCST_z3 | 1.4 (2) | 0.52 (4) | 0.69 (11) | 0.85 (15) | 0.69 (20) |
|  | 2080 | meanBLT_spwn, maxTLT_z1, maxTLT_z4, meanCST_mg | 1.73 (3) | 0.52 (3) | 0.69 (9) | 0.81 (58) | 0.69 (27) |
|  | 2072 | meanBLT_spwn, maxTLT_z1, maxTLT_z4, meanLST_z4 | 1.92 (4) | 0.51 (5) | 0.7 (7) | 0.84 (24) | 0.69 (32) |
|  | 2078 | meanBLT_spwn, maxTLT_z1, maxTLT_z4, meanCST_z4 | 1.94 (5) | 0.5 (8) | 0.7 (6) | 0.83 (33) | 0.69 (33) |
| Sorted by D(rand) | 2074 | meanBLT_spwn, maxTLT_z1, maxTLT_z4, meanLST_mg | 0.0 (1) | 0.55 (1) | 0.73 (1) | 0.86 (7) | 0.71 (10) |
|  | 4751 | meanBLT_spwn, maxTLT_z1, maxTLT_z4, meanBLT_jvnI0, meanLST_mg | 2.09 (6) | 0.52 (2) | 0.72 (4) | 0.87 (2) | 0.72 (1) |
|  | 2080 | meanBLT_spwn, maxTLT_z1, maxTLT_z4, meanCST_mg | 1.73 (3) | 0.52 (3) | 0.69 (9) | 0.81 (58) | 0.69 (27) |
|  | 2077 | meanBLT_spwn, maxTLT_z1, maxTLT_z4, meanCST_z3 | 1.4 (2) | 0.52 (4) | 0.69 (11) | 0.85 (15) | 0.69 (20) |
|  | 2072 | meanBLT_spwn, maxTLT_z1, maxTLT_z4, meanLST_z4 | 1.92 (4) | 0.51 (5) | 0.7 (7) | 0.84 (24) | 0.69 (32) |
| Sorted by D(block) | 2074 | meanBLT_spwn, maxTLT_z1, maxTLT_z4, meanLST_mg | 0.0 (1) | 0.55 (1) | 0.73 (1) | 0.86 (7) | 0.71 (10) |
|  | 4739 | meanBLT_spwn, maxTLT_z1, maxTLT_z4, maxTLT_mg, meanLST_mg | 2.66 (10) | 0.5 (10) | 0.72 (2) | 0.86 (11) | 0.71 (4) |
|  | 4758 | meanBLT_spwn, maxTLT_z1, maxTLT_z4, meanLST_z1, meanLST_mg | 2.62 (9) | 0.48 (19) | 0.72 (3) | 0.84 (17) | 0.71 (3) |
|  | 4751 | meanBLT_spwn, maxTLT_z1, maxTLT_z4, meanBLT_jvnI0, meanLST_mg | 2.09 (6) | 0.52 (2) | 0.72 (4) | 0.87 (2) | 0.72 (1) |
|  | 4767 | meanBLT_spwn, maxTLT_z1, maxTLT_z4, meanLST_mg, meanCST_z1 | 3.1 (17) | 0.47 (25) | 0.71 (5) | 0.86 (8) | 0.71 (8) |
| Sorted by D(pred) | 4750 | meanBLT_spwn, maxTLT_z1, maxTLT_z4, meanBLT_jvnI0, meanLST_z5 | 4.08 (27) | 0.48 (23) | 0.67 (24) | 0.87 (1) | 0.7 (16) |
|  | 4751 | meanBLT_spwn, maxTLT_z1, maxTLT_z4, meanBLT_jvnI0, meanLST_mg | 2.09 (6) | 0.52 (2) | 0.72 (4) | 0.87 (2) | 0.72 (1) |
|  | 4754 | meanBLT_spwn, maxTLT_z1, maxTLT_z4, meanBLT_jvnI0, meanCST_z3 | 2.98 (13) | 0.49 (16) | 0.66 (34) | 0.87 (3) | 0.71 (6) |
|  | 4768 | meanBLT_spwn, maxTLT_z1, maxTLT_z4, meanLST_mg, meanCST_z2 | 2.4 (7) | 0.49 (13) | 0.69 (8) | 0.87 (4) | 0.72 (2) |
|  | 4748 | meanBLT_spwn, maxTLT_z1, maxTLT_z4, meanBLT_jvnI0, meanLST_z3 | 4.14 (28) | 0.46 (43) | 0.66 (38) | 0.86 (5) | 0.7 (17) |
| Sorted by D(total) | 4751 | meanBLT_spwn, maxTLT_z1, maxTLT_z4, meanBLT_jvnI0, meanLST_mg | 2.09 (6) | 0.52 (2) | 0.72 (4) | 0.87 (2) | 0.72 (1) |
|  | 4768 | meanBLT_spwn, maxTLT_z1, maxTLT_z4, meanLST_mg, meanCST_z2 | 2.4 (7) | 0.49 (13) | 0.69 (8) | 0.87 (4) | 0.72 (2) |
|  | 4758 | meanBLT_spwn, maxTLT_z1, maxTLT_z4, meanLST_z1, meanLST_mg | 2.62 (9) | 0.48 (19) | 0.72 (3) | 0.84 (17) | 0.71 (3) |
|  | 4739 | meanBLT_spwn, maxTLT_z1, maxTLT_z4, maxTLT_mg, meanLST_mg | 2.66 (10) | 0.5 (10) | 0.72 (2) | 0.86 (11) | 0.71 (4) |
|  | 3407 | meanBLT_precond, meanBLT_spwn, maxTLT_z1, maxTLT_z4, meanLST_mg | 2.96 (12) | 0.5 (9) | 0.61 (146) | 0.86 (10) | 0.71 (5) |

The top ranked model according to ΔAICc for Washington, Oregon, and Northern California, differed from those ranked best according to D(rand), D(block), D(pred) or D(total). In most cases the best model according to ΔAICc was not one of the five top ranked models by other metrics (Tables 2–4). The top five models for Washington, when ranked by ΔAICc had a ΔAICc<2. Those same five models had a lower D(rand) (0.25), and D(block) (0.17) when compared with Oregon and Northern and Central California. The top ranked models for Oregon when sorted by D(rand), D(block), and D(pred), led to metric values of 0.34, 0.54 and 0.53 respectively. The corresponding values for Northern California were 0.31, 0.50 and 0.70 (Table 4).

### D(block) as a metric to select ‘best’ model

We used D(block) to select the ‘best’ models because it measures the performance of models while accounting for temporal structure (e.g., autocorrelation) in the data. Moreover, this metric has been shown to outperform other approaches when predictive accuracy is sought (e.g., Roberts et al., 2017; Kiaer et al., 2021). The resulting ‘best’ models were therefore 12295 (Washington), 1853 (Oregon), 1602 (Northern California), and 2074 (Central California). The ‘best’ model according to D(block) was also the top ranked model according to ΔAICc, and D(rand) in Central California. For Washington, Oregon, and Northern California, the ‘best’ model according to D(block) differed from the top ranked models according to the other metrics.

Adding two-year and one-year lagged responses as independent variables did not improve model performance, especially when model performance was based on our metric of choice, D(block) (Tables S14-S17). The top performing models according to D(block) did not include any lagged response variables and out-performed models with lagged response variables for all regions. Moreover, the top ranked models according to D(block) were the same irrespective of whether the one-year and two-year lagged responses were included as possible covariates. For instance, in Central California, the top models according to D(block) were the same when the lagged response is considered as a possible covariate even though the number of data points is two fewer for the analyses that explored lags (Tables 5 and S8).

### Common environmental processes among top ranked models

In general, each region had different variables that appeared most often in the top five models and the selected variables differed among model selection methods. There was more commonality in the selected independent variables for Washington and Central California (Tables 2 and 5). Average temperature of the bottom water layer during females preconditioning (meanBLT_precond) and the maximum temperature of the top water layer during the zoeal stage 3 (maxTLT_z3) were the independent variables selected most consistently for Washington according to ΔAICc, D(rand), D(block), and D(total). In Central California, the average temperature of the bottom water layer during spawning (meanBLT_spwn) and the maximum temperature of the top water layer during zoeal 1 and zoeal 4 (maxTLT_z1 and maxTLT_z4) were the most common independent variables in the top ranked models according to the five metrics.

There was commonality among the models selected by only some of the metrics for Oregon and Northern California (Tables 3 and 4). For example, in Oregon, the average cross-shelf transport of the top water layer during zoeal 4 (meanCST_z4) was the independent variable selected most consistently according to ΔAICc, D(rand), D(block), D(pred), and D(total). Additionally, average long-shore transport and the average temperature of the bottom water layer during juvenile age-0 (meanBLT_jvnl0) was the independent variable selected most consistently according to ΔAICc, D(rand), and D(total). The average temperature of the top water layer during zoeal 3 (meanTLT_z3) and the average long-shore transport of the top water layer during zoeal 5 (meanCST_z5) were the independent variables selected most consistently according to ΔAICc, D(rand), and D(total) for Northern California.

An initial analysis of correlations between pre-season abundances from the four regions found that preseason abundance of Oregon Dungeness crab explained 60% of the variation in preseason abundance of Dungeness crab off Northern California (Fig. S10). In contrast, the correlations between Central/Northern California (R^2^=12.51%) and Oregon/Washington (R^2^=2.36%) were much lower. Two hypotheses can explain the high correlation between Northern California and Oregon: 1) the two populations are driven by similar oceanographic factors, or 2) there is significant larval transport from one population to another. However, we could not find any oceanographic drivers for both populations to support the first hypothesis (Tables 2–5). To test the latter hypothesis, we included transport variables from one region as independent variables in the other regions and reran the model selection process. The latter analysis was conducted because if there were a larval transport from Oregon to Northern California, we would expect long-shore transport variables to be selected in the best models and that the resulting models would perform better that the model identified above for Northern California.

The inclusion of transport variables from Northern California as predictors of pre-season abundance of Oregon Dungeness crab did not improve the performance of the ‘best’ model according to D(block) (Table S18). However, the inclusion of transport variables from Oregon as predictors of pre-season abundance of Dungeness crab off Northern California improved the performance of ‘best’ model according to D(block), D(pred), and D(total) (Table S19). D(block), D(pred), and D(total) increased from 0.5, 0.75, and 0.62 to 0.75, 0.82, and 0.67, respectively. Moreover, best model according to D(block) had long-shore transport variables during zoeal 2. This result is evidence of larval transport from Oregon to Northern California, especially during early larval stages, which in turn contributed to the size of the fishable population four years later.

### The effects of oceanographic variables on preseason abundances

Analysis of partial residuals and the standardized coefficients of regression of the ‘best’ models can help explain the mechanisms through which oceanographic variables affect preseason abundance. We standardized the coefficients of regression by dividing them by their standard deviation. The results (Table 6) indicate that each region differs in terms of mechanisms that affect pre-season abundance. The standardized regression coefficients suggest greatest effects of the maximum temperature of the top water layer during zoeal 3 in Washington, the average long-shore transport of the top water layer during zoeal stage 1 in Oregon, average temperature of the top water layer during zoeal stage 5 in Northern California, and average temperature of the bottom water layer during spawning in Central California.

**Table 6:** Standardized coefficients of regression and variance inflation factors (VIF) for ‘best’ model per region.

|  | Variable/model | Estimate | Std. Error | Pr(>\|t\|) | VIF |
| --- | --- | --- | --- | --- | --- |
| **WA** | (Intercept) | 2.0445 | 0.0665 | 0.0000 |  |
|  | meanBLT_precond | 0.1686 | 0.0825 | 0.0521 | 1.4879 |
|  | maxTLT_z3 | -0.2649 | 0.0819 | 0.0035 | 1.4648 |
|  | maxTLT_z5 | 0.0839 | 0.0795 | 0.3019 | 1.3815 |
|  | maxTLT_mg | 0.1107 | 0.0786 | 0.1717 | 1.3490 |
|  | meanLST_z3 | -0.1582 | 0.0806 | 0.0613 | 1.4195 |
| **OR** | (Intercept) | 1.9145 | 0.0651 | 0.0000 |  |
|  | meanBLT_precond | -0.1362 | 0.0780 | 0.0930 | 1.3849 |
|  | meanBLT_spwn | -0.1975 | 0.0770 | 0.0167 | 1.3512 |
|  | meanLST_z1 | 0.2965 | 0.0794 | 0.0009 | 1.4368 |
|  | meanCST_z4 | -0.1840 | 0.0807 | 0.0314 | 1.4837 |
| **N.CA** | (Intercept) | 1.3430 | 0.0901 | 0.0000 |  |
|  | maxTLT_z3 | -0.1598 | 0.0942 | 0.1015 | 1.0546 |
|  | meanTLT_z5 | -0.3255 | 0.1039 | 0.0042 | 1.2853 |
|  | meanLST_z3 | 0.1176 | 0.1015 | 0.2569 | 1.2255 |
| **C.CA** | (Intercept) | 0.4034 | 0.0935 | 0.0002 |  |
|  | meanBLT_spwn | 0.6928 | 0.1372 | 0.0000 | 2.0795 |
|  | maxTLT_z1 | -0.4010 | 0.1668 | 0.0239 | 3.0716 |
|  | maxTLT_z4 | -0.5633 | 0.1593 | 0.0016 | 2.8025 |
|  | meanLST_mg | -0.3125 | 0.1121 | 0.0100 | 1.3886 |

### Sensitivity of ‘best’ models to individual years

Removing individual years and refitting the ‘best’ models (jackknifing) had little impact on the model fit in the four regions (Figs S15-S18), the median D(total) over the remaining 29 years for Washington, Oregon, and Northern and Central California were 0.36, 0.52, 0.38 and 0.70 respectively. Moreover, predicting the missing year from any iteration produced estimates very similar to those for the full data set (blue points in the Fig. 3). In Washington, the years that exhibited the highest impact on the ability to explain the data were 1992 (D(total) increased from 0.36 to 0.45), and 1989 (D(total) decreased to 0.29). In Oregon, the most influential years were 1998 (D(total) increased from 0.52 to 0.58) and 2006 (D(total) decreased to 0.47). In Northern California, the most influential years were 1996 (D(total) increased from 0.38 to 0.52) and 2002 (D(total) decreased to 0.31). In Central California, the most influential years were 2010 (D(total) increased from 0.70 to 0.73) and 2012 (D(total) decreased to 0.66).

**Figure 3:**
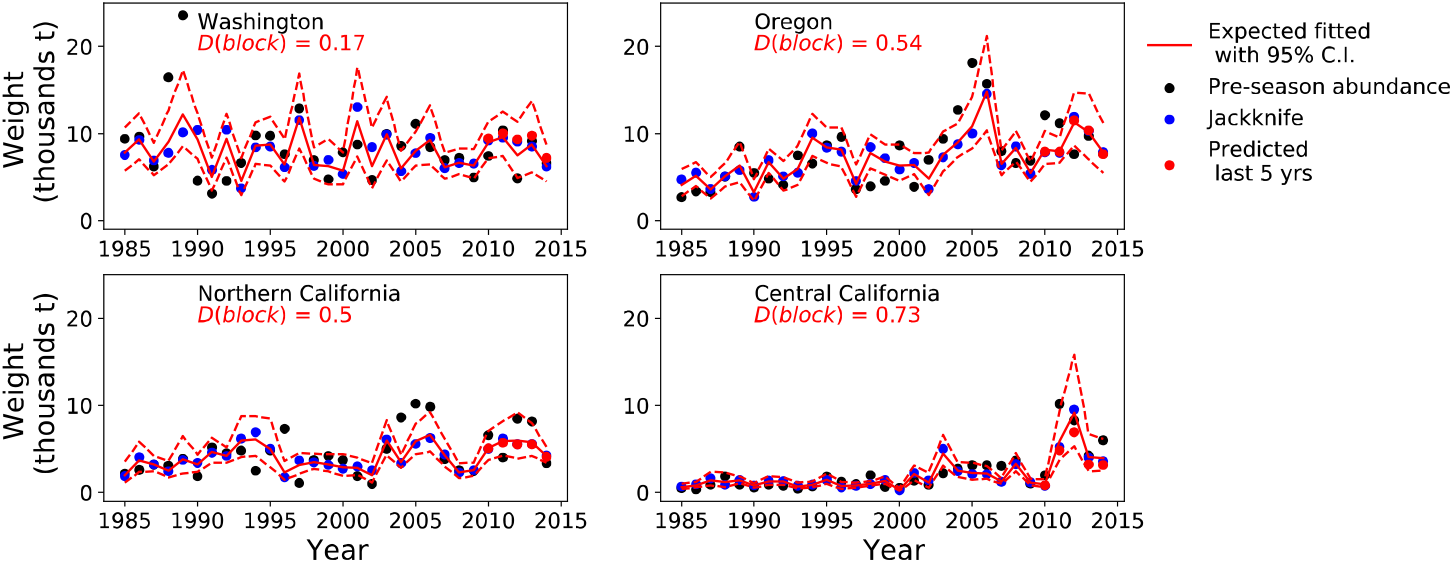
Fit of the ‘best’ models to pre-season abundance for Washington, Oregon, and Northern and Central California. The solid red line is the predicted abundance with 95% confidence intervals from the full time series. Dark points are the estimates of the preseason abundance of legal-sized male Dungeness crab from Richerson et al. (2020). The blue points are predicted values from jackknife analysis removing individual years one data point at a time. Red points are predictions from fitting the best-fit model to data 1981-2009 and then predicting 2010-2014.

## Discussion

We started with the idea of developing an environmental index for forecasting crab abundance in advance of the fishery. This goal requires building a model that not only can explain existing data, but also can accurately predict out of sample outcomes. We used five metrics for model selection, each of which differs in how it defines model performance and skill on available and out of sample data. Model selection was generally not robust to the metric used, with each metric often resulting in different model rankings, illustrating the need to carefully choose performance metrics for model selection. The ‘best’ models following D(block) suggested that the mechanisms that drive pre-season abundance vary among regions.

While the ‘best’ models following D (block) were selected for predictive accuracy, in our case they also often had high explanatory accuracy. The ‘best’ models according to D (block) explained 36%, 53%, 38% and 71% of the variation according to D(total) for the whole time series of preseason abundances in Washington, Oregon and Northern and Central California, respectively. This implies that the best models according to D(block) can capture the relationships between oceanographic variables with preseason abundance, hence implicitly the mechanisms that drive preseason abundance. Moreover, the ‘best’ models approximately (because the metrics are not based on conventional *R*^2^ statistics, except D(total)) satisfy the guideline by de Oliveira and Butterworth (2005) that at least 50% of the total variation in recruitment needs to be explained for an environment index to be meaningful for management in Oregon and Central California.

### Model selection metric

Unlike other studies that used a single metric for model selection (e.g., Johnson & Omland, 2004), we used five metrics. The five metrics were based on cross validation, corrected Akaike Information Criterion (AICc), and explained variance. AIC is a popular method of comparing multiple models while considering explanatory accuracy and parsimony (Anderson & Burnham, 2004). Cross-validation methods, on the other hand, select models according to their predictive accuracy. D(rand), D(block), and D(pred) are three examples of cross-validation methods, with the last two accounting for correlation among data. Although D(rand) is computationally expensive, it may be more appropriate when the dependent variable does not exhibit temporal or spatial autocorrelation. However, autocorrelation is typical for ecological data sets (Pimm et al., 1988). D(pred) is more appropriate for data with a constant trend that does not change with time, but predictive accuracy for recent years may not hold in future years if the processes driving recruitment change over time (e.g., due to climate cycles or regime shifts (Deyoung et al., 2008). Compared to the other metrics we used, we determined D(block) to be most appropriate metric for model selection given that it can achieve the two goals of evaluating the ability to explain past data and prediction skill and because it accounts for the temporal structure of the data.

The differences in model performance among the five metrics suggests that models might have a higher explanatory accuracy for past data but a low predictive accuracy and vice-versa, and the differences among the models selected using D(rand), D(block), and D(pred) suggest that predictive accuracy depends on the structure of the data (Roberts et al., 2017). Models that fit the data well or predict random holdout samples may provide poor predictions in recent years and vice versa. These differences between rankings of models among the five metrics indicate that the models selected using D(block) still depend on the data used when selecting models and may not perform as expected on other (future) data. For example, the best model according to D(block) for Washington did not perform well for the most recent years (Table 2). Therefore, it is useful to use multiple metrics that can identify strengths and limitations of alternative models when selecting models for predictive accuracy. It may be useful to consider how predictions are likely to be used, and the implications of predictive accuracy on the value provided by predictions when choosing a metric for model selection (Kiaer et al., 2021).

### Implications for drivers of Dungeness crab recruitment

The results suggest that the performances of the best models and the mechanisms that drive Dungeness crab preseason abundance appear to differ among regions despite similar regional life histories. The life-stage- and spatiotemporally-specific mechanistic hypotheses considered in the analyses cover many of the processes that can affect survival and recruitment, including horizontal transport. Compared to other studies that suggest horizontal transport as the only mechanism that drives the abundance of megalopae (Shanks, 2013), our results suggest that female preconditioning and spawning conditions are drivers of pre-season abundance for Dungeness crab off the US West coast.

Temperature can have either a positive or a negative effect depending on life stage, with generally positive effects during female precondition and spawning stages for Washington and Central California, negative effects during zoeal 3 (Washington) and zoeal 1 and 4 (Central California), and positive effects during the zoeal 5 and megalopae stages (Washington). Similarly, transport variables, especially long-shore transport, have either a positive effect on the abundance during the early larval stages, such as zoeal 1 for Oregon and zoeal 3 for Northern California, or a negative effect during the late larval stages such as during the megalopae for Central California.

Correlation does not prove causation. However, we can make several hypotheses regarding relationships between temperature and recruitment. Given adequate food resources, warmer temperature can allow for faster growth and larger female size, which may result in higher fecundity (Shields, 2020). The positive effects of temperature can extend beyond female preconditioning to the larval phases by helping larvae to grow faster, leading to lower vulnerability to size-based predation (Sulkin & McKeen, 1989). This may be the case for Washington where we found a positive effect of temperature during the female preconditioning, zoeal stage 5 and the megalopae stage. Additionally, while spawning success may decline at higher temperature, (Mayer, 1973; Wild, 1983b) increased egg growth and fecundity in warmer water may explain why we found a positive effect of temperature during the spawning phase for Washington and Central California. Temperature can also have a negative effect especially during early larval stages when individuals are most vulnerable to reduced growth and high mortality when temperature exceeds optimal limits (Sulkin et al., 1996). This effect may be the case for Central California where we found a negative effect of temperature on the zoeal 1 and 4 stages. Additionally, warmer temperatures increase metabolism, which may result in less energy for reproduction during the female preconditioning stage or increased risk of starvation for feeding larvae if food resources are not adequate.

Although the transport-related variables appeared in the ‘best’ models for all four regions, their contribution was less than that of temperature-related variables, especially for Washington, Northern and Central California. In Oregon, northward transport of larvae during zoeal 1 positively affected preseason abundance. Other studies suggest that megalopae may originate from outside Oregon during high abundance years (Shanks, 2013). The ‘best’ model in Oregon followed trends in pre-season abundance, particularly during the years 2007-2012 when Shanks (2013) reported extremely high recruitment of megalopae, but pre-season abundance during the same years appeared to be slightly above the historical average. This suggests that the high recruitment of megalopae may be specific to the Charleston, Oregon marina. However, it is plausible that there may be larval transport between regions, especially from Oregon to Northern California.

Climate change will drive changes in both temperature and transport. It is difficult to predict how warming temperatures will influence Dungeness crab based on our modeling; our models were not able to estimate non-linear relationships between pre-season abundance and temperature-related variables, even though survival must decline at high enough temperatures based on fundamental bioenergetics. Transport-related variables will play a crucial future role, especially for Oregon and Northern California, where transport positively affects Oregon pre-season abundance. This result is consistent with the suggestion that transport will play a crucial role for species with longer (>3 months) larval durations such as Dungeness Crab (Bani et al., 2020).

The question of whether the Dungeness crab pre-season abundance is controlled by processes before or after settlement to the benthos at age-0 is unknown. Our study did not include processes for juveniles between age-0 and age-4, when Dungeness crab are of legal size. However, our resulting ‘best’ model followed the trends of the estimated pre-season abundance for all four regions (Fig. 3). In Washington, and Northern and Central California, temperature-related factors are the most important post-settlement processes we found that affect pre-season abundance, while the transport-related variables are the most important processes we found for Oregon.

## Conclusion

It is important to understand the potential pitfalls and benefits associated with alternative model selection criteria given that model selection results are sensitive to the criterion applied. Future studies should both consider multiple model selection criteria as well as clearly state the objectives for the study (e.g., understanding processes governing recruitment, making predictions). We focused on models selected using D(block), a metric designed to balance explanatory and predictive ability. We found that the processes determining pre-season abundance could be identified with sufficient precision to enable a predictive skill, suggesting that the predictions may be useful for management purposes. Moreover, we found that transport (within and between regions), as well as temperature were likely drivers of pre-season abundance, highlighting that future studies should focus on multiple processes, ideally guided by an initial identification such as Table 1.

## Acknowledgments

This work was funded by the National science Foundation grant no. 1616821.

## Data availability Statement

Code to process the data and analysis are available at [www.github.com/ridouanbani].

