## Supplementary figures and materials for "Oceanographic drivers of legal-sized male Dungeness crab in the California Current System"

**Figure S1:** The spatial scale over which the independent variables are calculated. The top left figure shows the spatial scale (shallow waters, 0-8km offshore) over which the spawning phase is assumed to occur. Top right figure shows the spatial (continental shelf) scale over which the female preconditioning, and the zoeal 1 and 2 stages are assumed to occur. The bottom left figure shows the spatial scale (continental shelf and up to 150km offshore) over which the zoeal 3 and 4 stages are assumed to occur. The bottom right figure shows the spatial scale (continental shelf and up to 250 km offshore) over which the zoeal 5 and megalopae stages are assumed to occur.

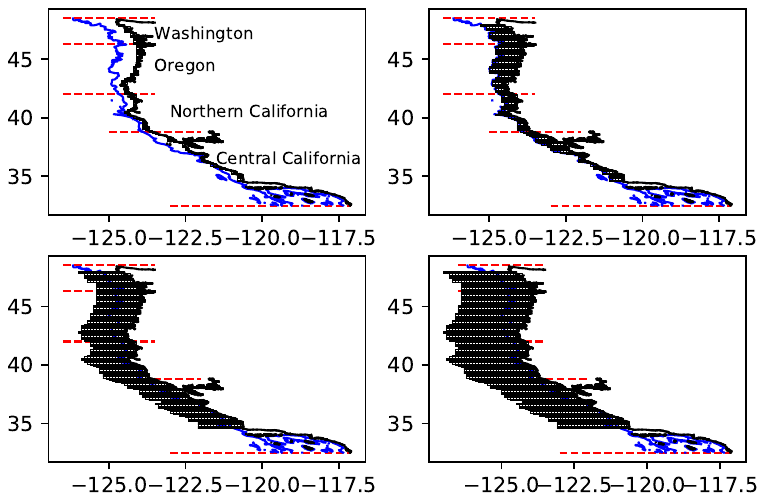

**Figure S2:** Average temperature of the water top layer (mean_TLT) during zoeal 1, 2, 3, 4, and 5 and megalopae in the four regions of Washington, Oregon, Northern and Central California

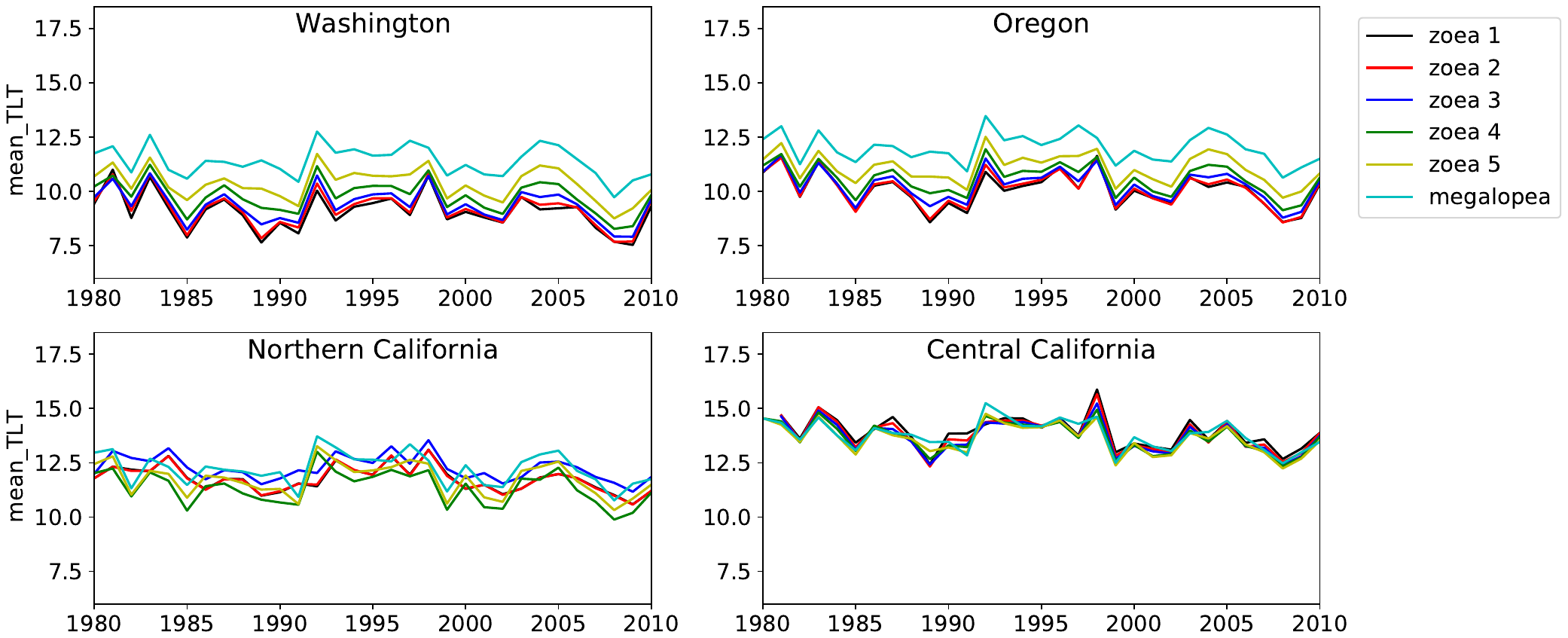

**Figure S3:** Maximum temperature of the water top layer (max_TLT) during zoeal 1, 2, 3, 4, and 5 and megalopae in the four regions of Washington, Oregon, Northern and Central California

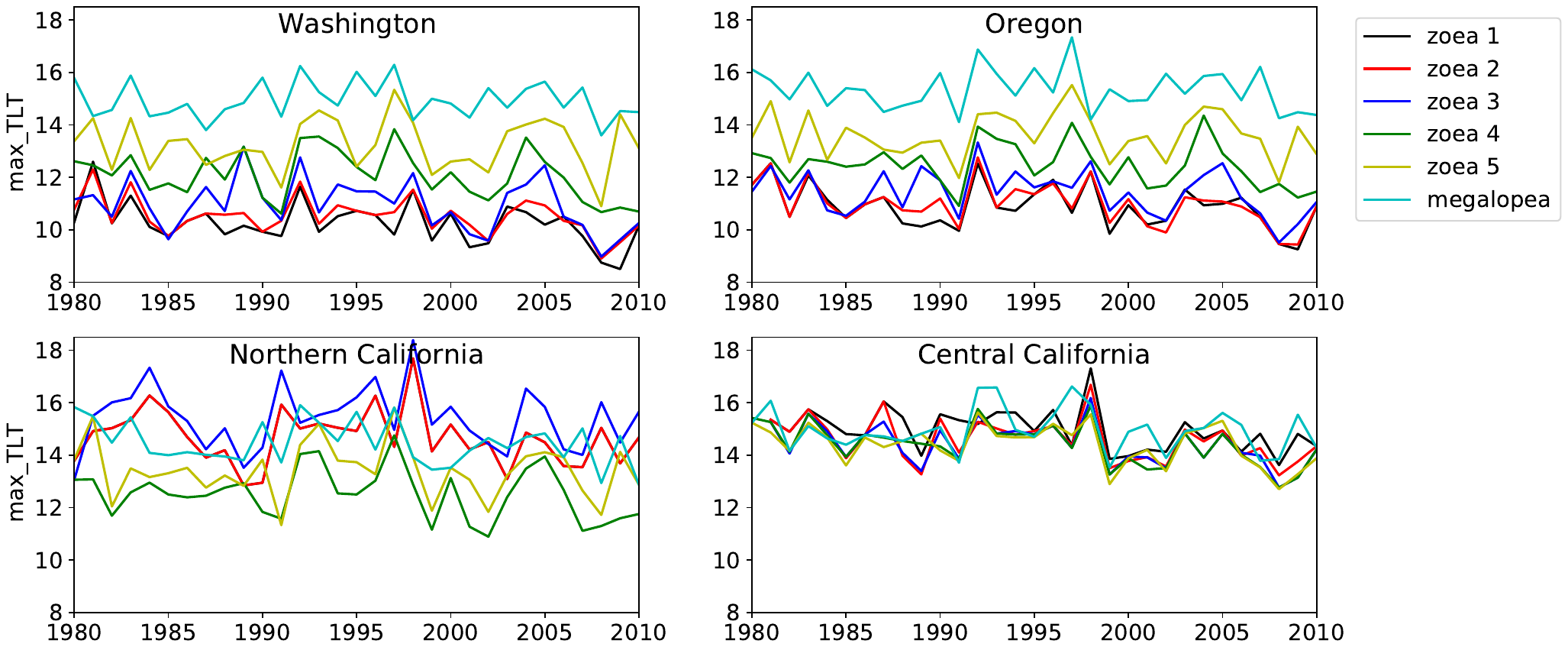

**Figure S4:** Average cross-shelf transport of the water top layer (mean_CST) during zoeal 1, 2, 3, 4, and 5 and megalopae in the four regions of Washington, Oregon, Northern and Central California

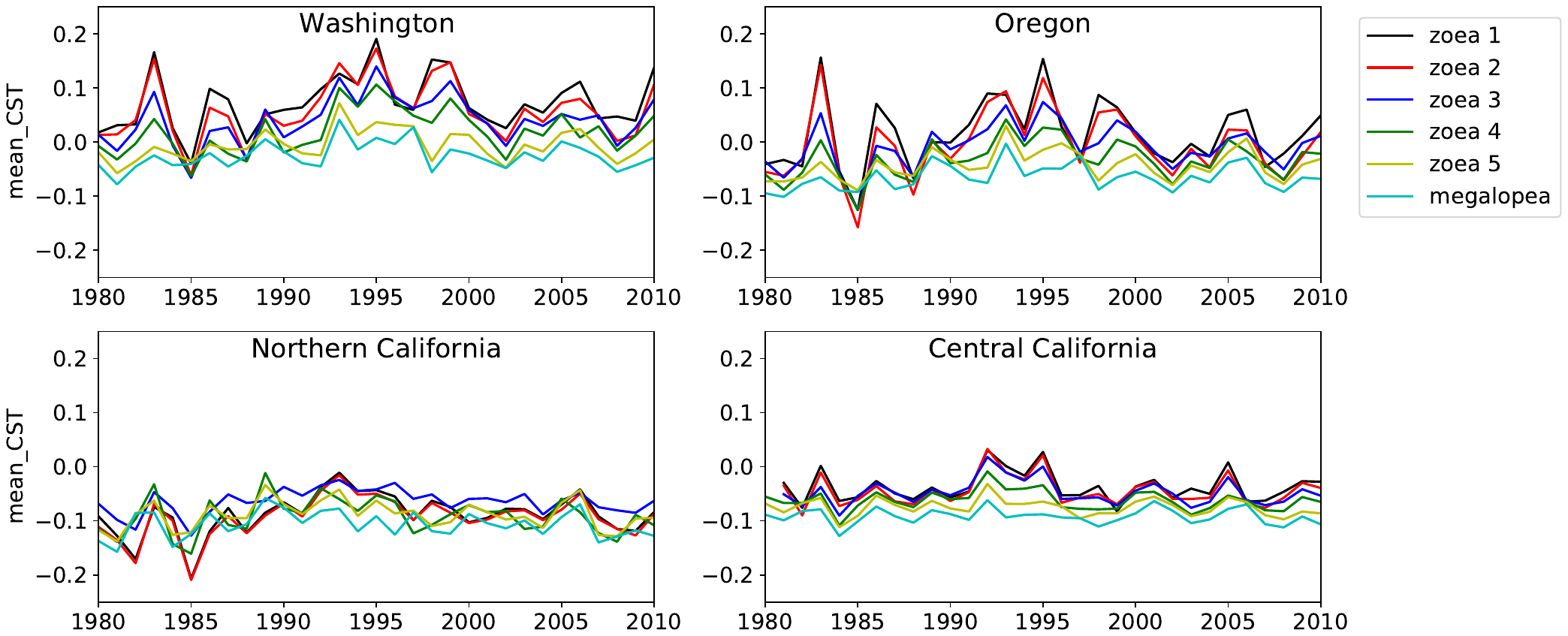

**Figure S5:** Average long-shore transport of the water top layer (mean_LST) during Zoeal 1, 2, 3, 4, and 5 and megalopae in the four regions of Washington, Oregon, Northern and Central California

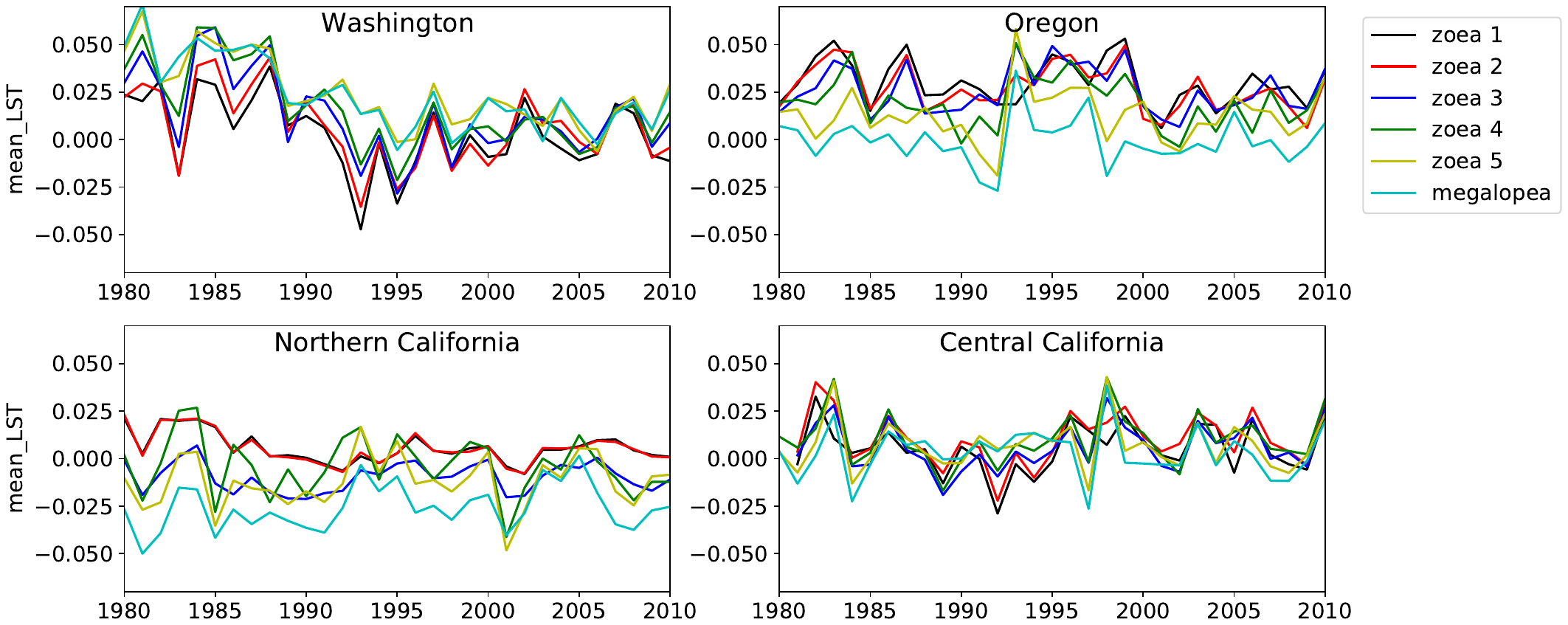

**Table S1:** Correlations among independent variables for Washington

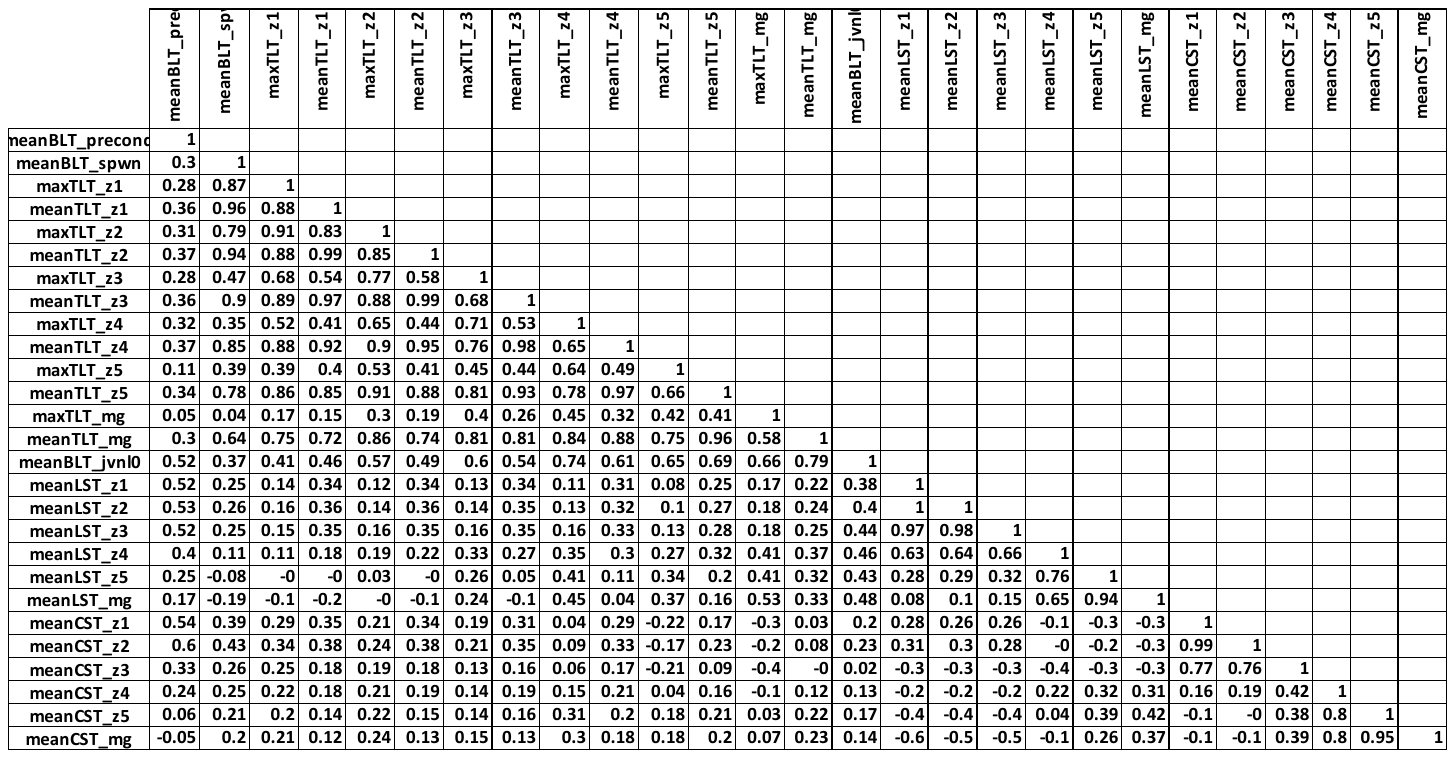

**Table S2:** Correlations among independent variables for Oregon

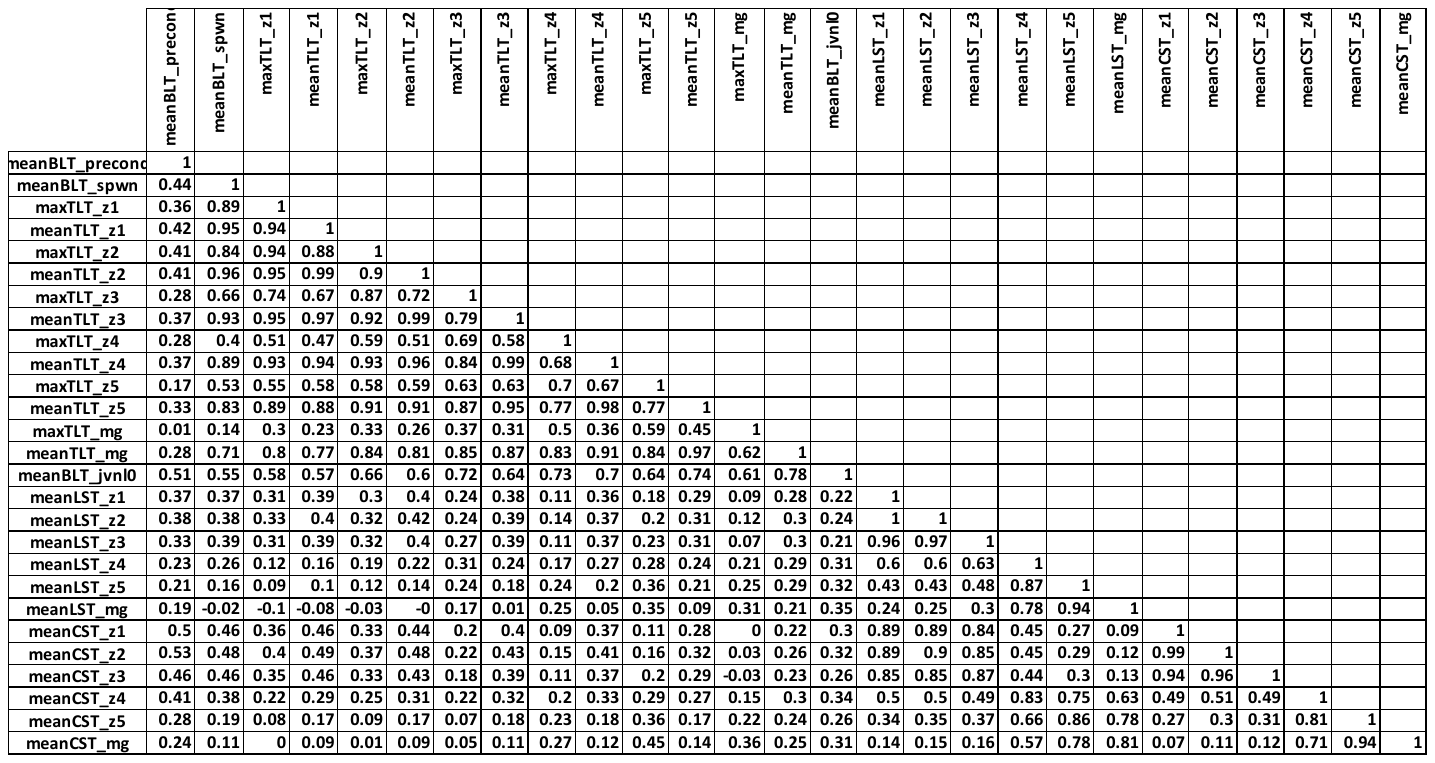

**Table S3:** Correlations among independent variables for Northern California

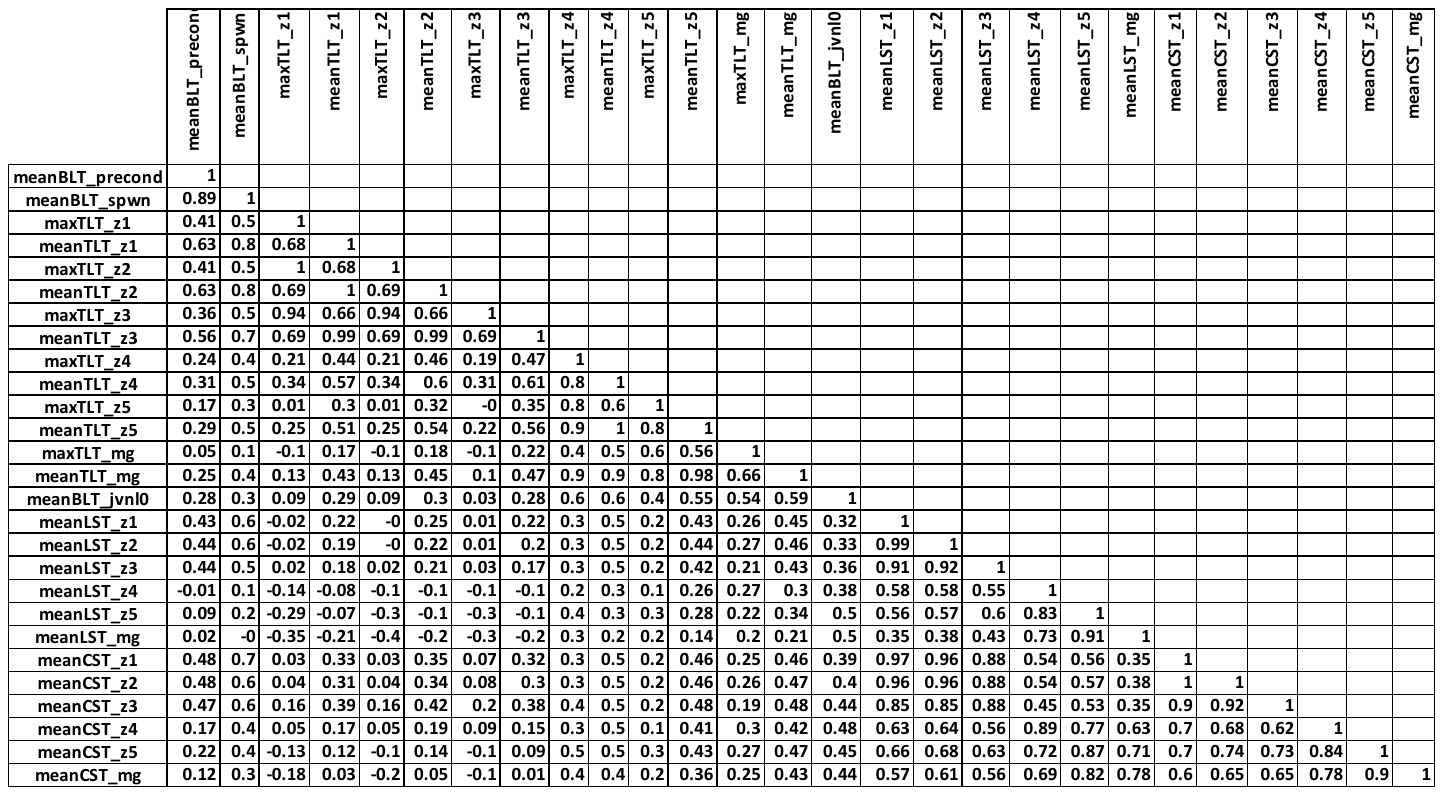

**Table S4:** Correlations among independent variables for Central California

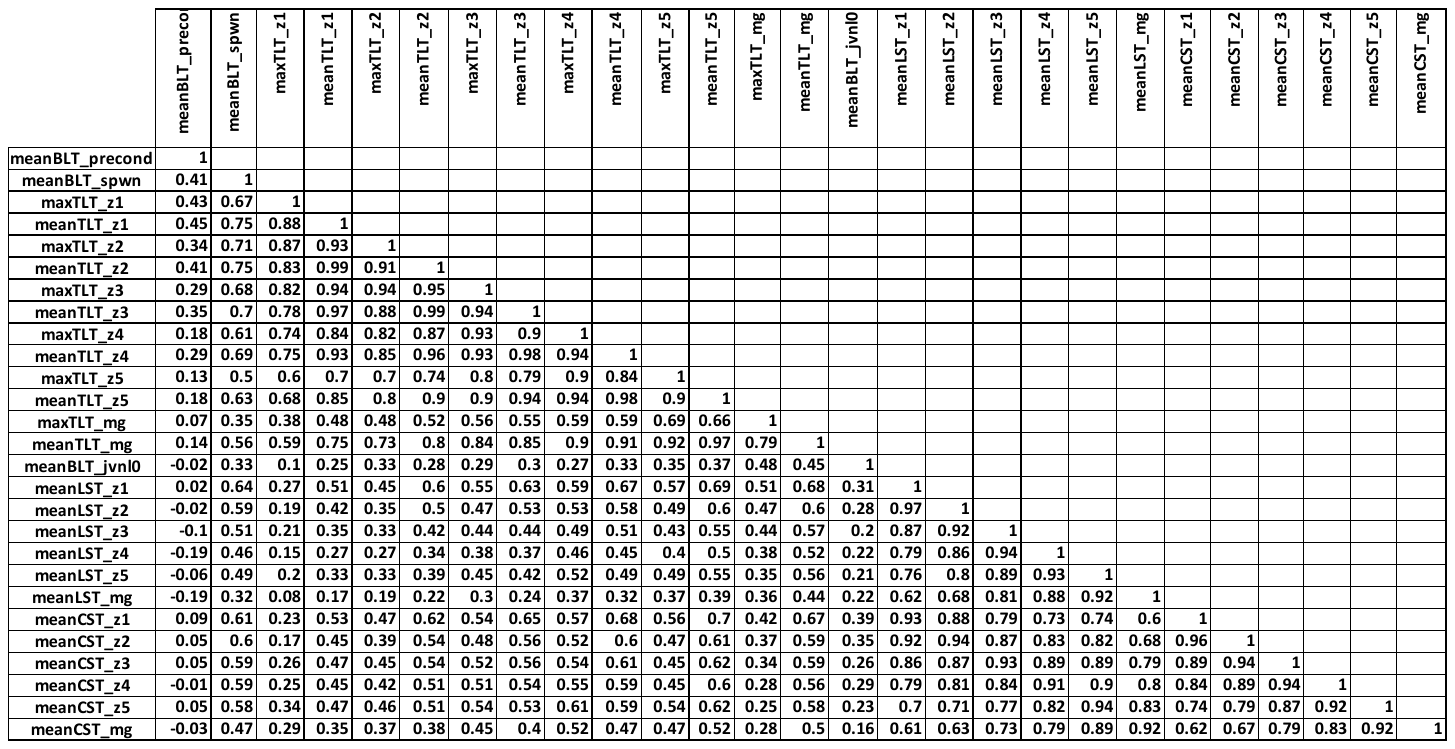

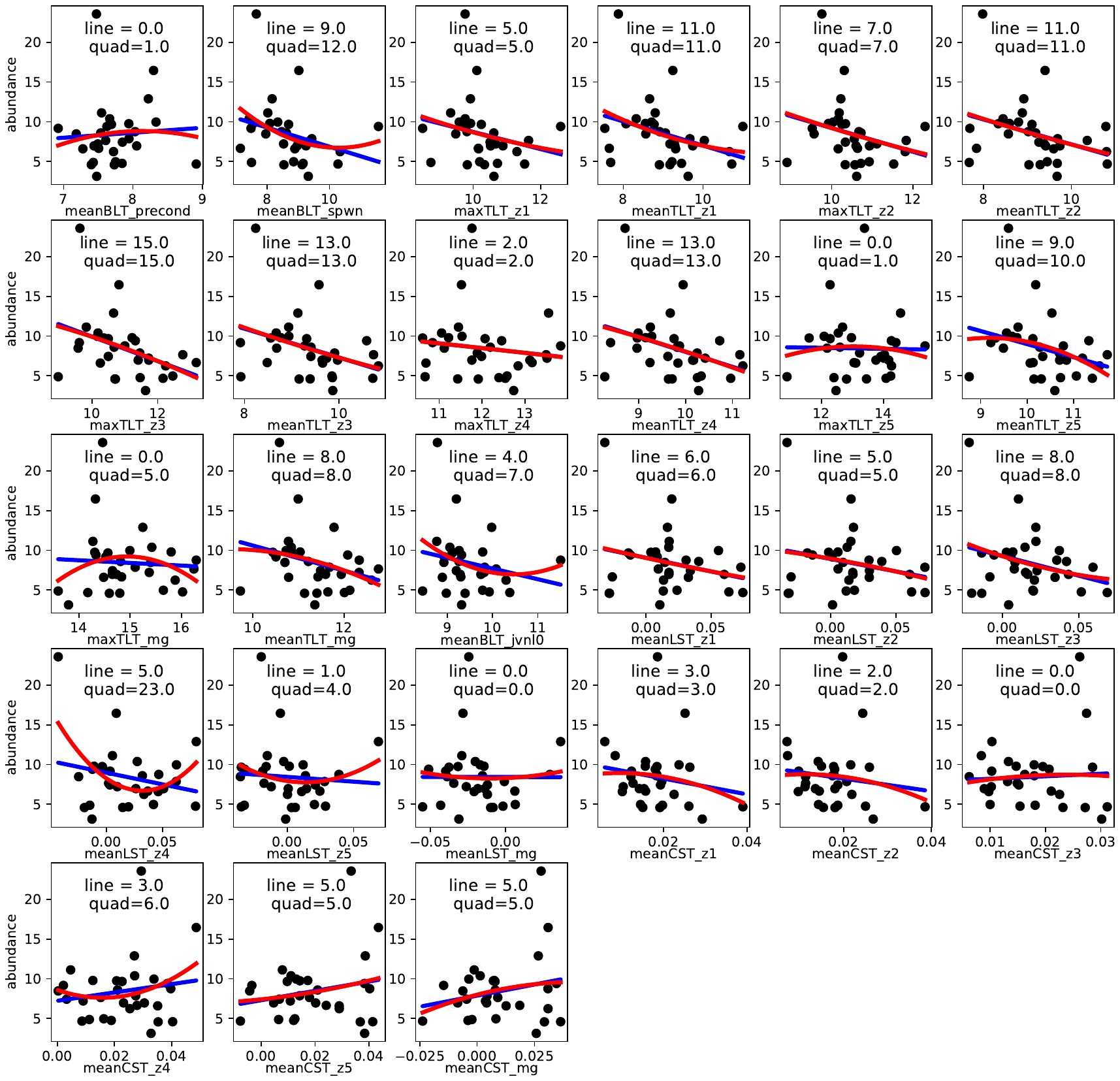
**Figure S6**: Linear (blue) v.s. quadratic (red) fit of the preseason abundance as a function of independent variables for Washington. The *R*^2^ values for the line and quadratic functions provided within each panel.

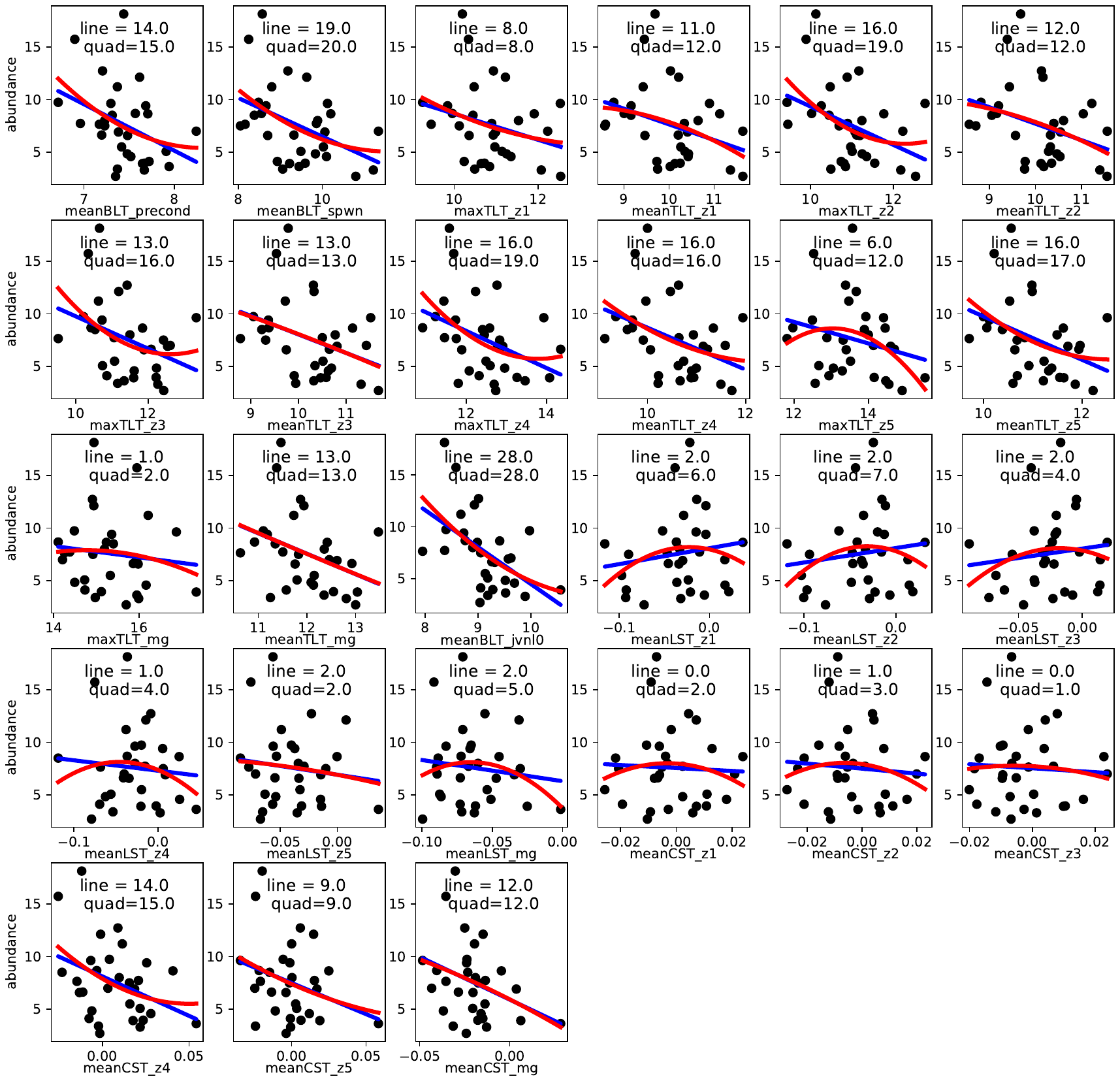
**Figure S7**: Linear (blue) v.s. quadratic (red) fit of the preseason abundance as a function of independent variables for Oregon. The *R*^2^ values for the line and quadratic functions provided within each panel.

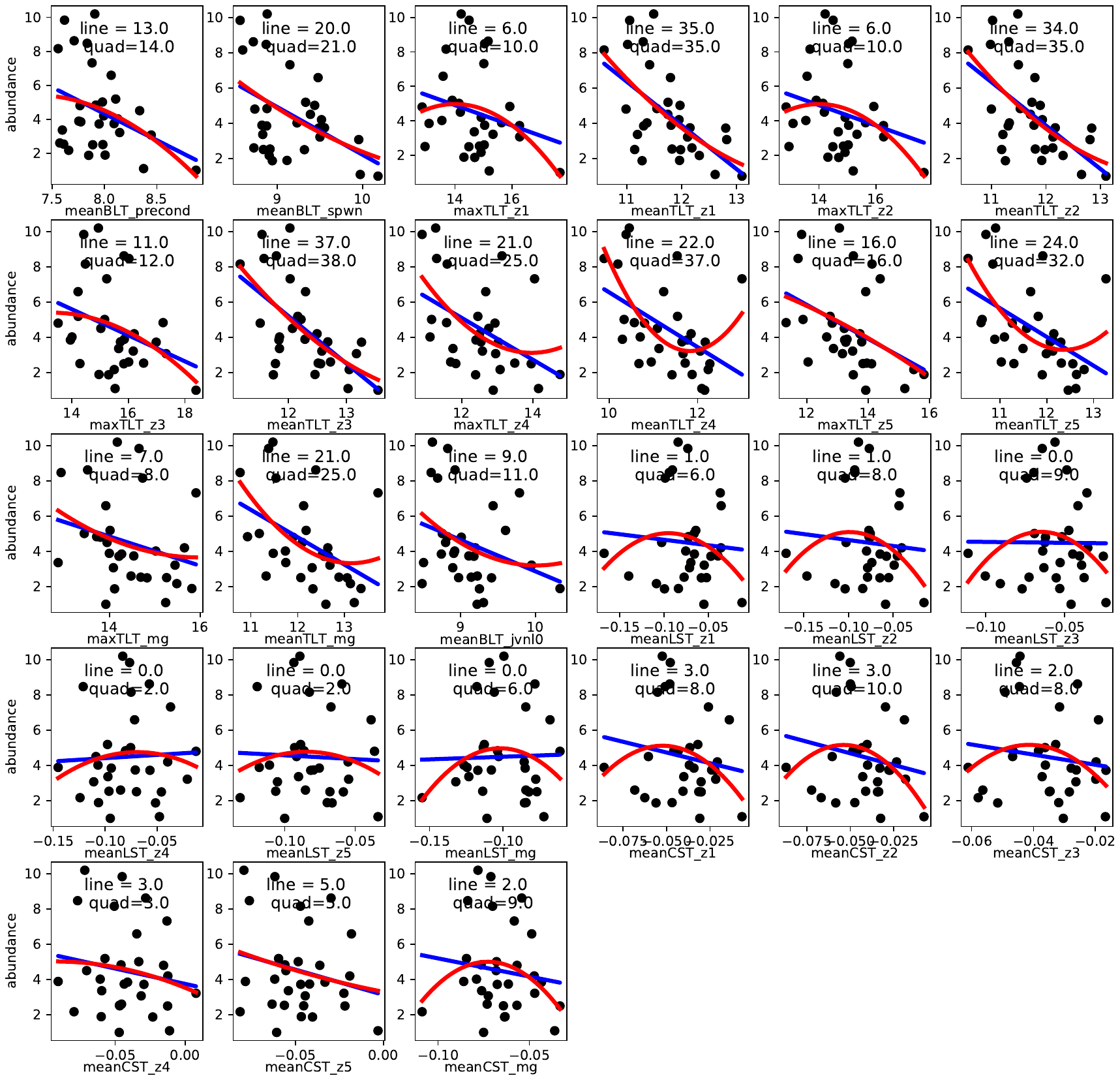
**Figure S8**: Linear (blue) v.s. quadratic (red) fit of the preseason abundance as a function of independent variables for Northern California. The *R*^2^ values for the line and quadratic functions provided within each panel.

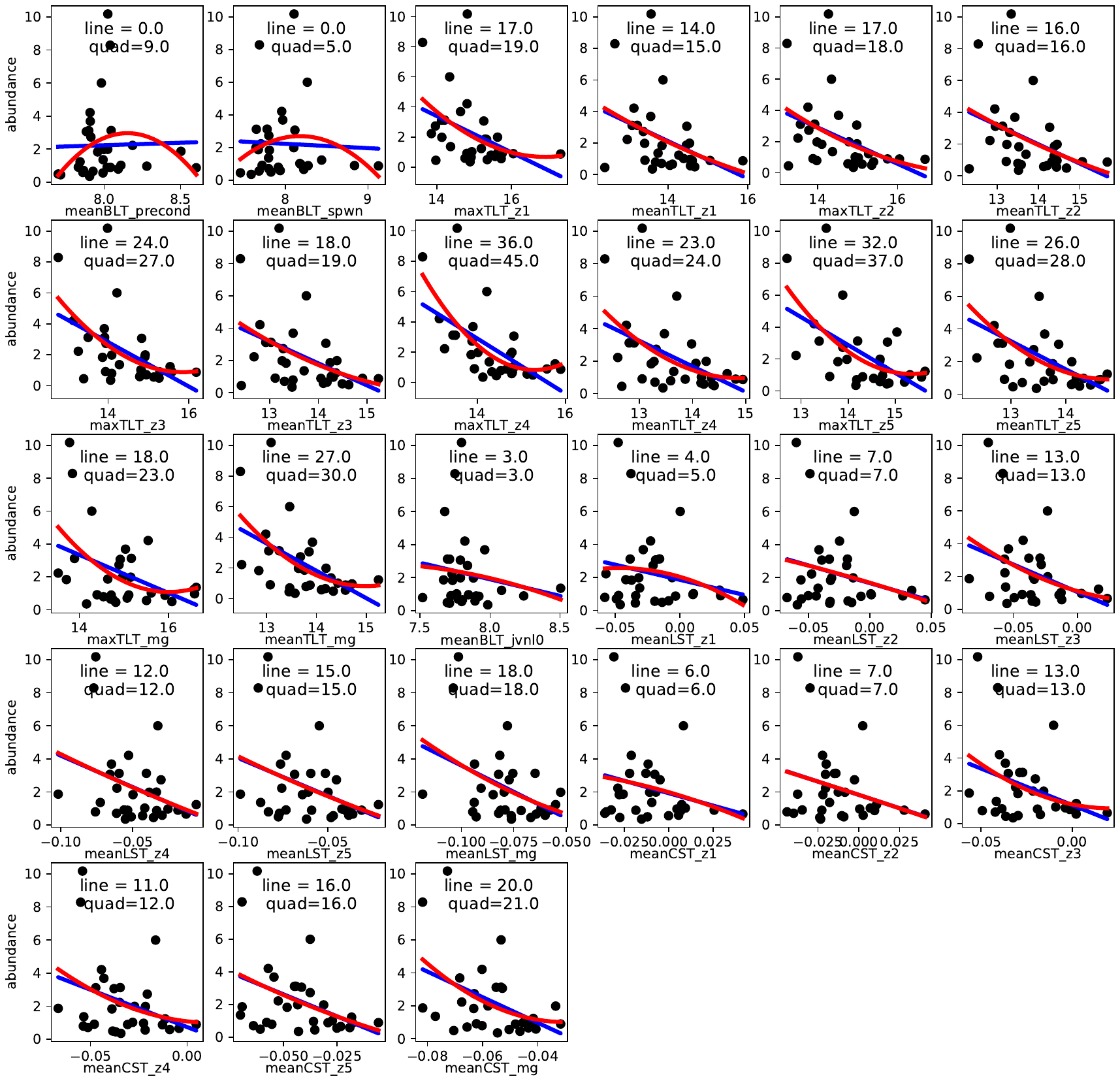
**Figure S9**: Linear (blue) v.s. quadratic (red) fit of the preseason abundance as a function of independent variables for Central California. The *R*^2^ values for the line and quadratic functions provided within each panel.

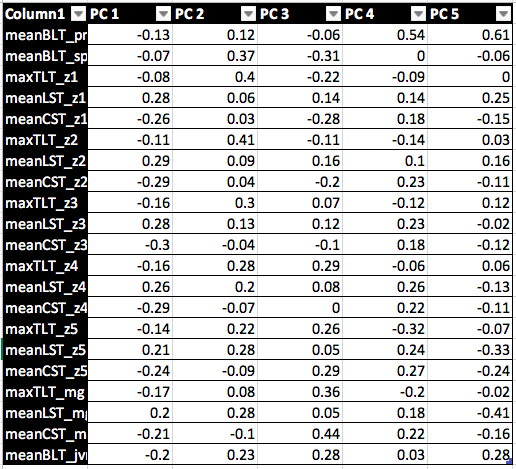

**Table S5**: Table showing the five principal components for Washington and their loadings in term of ROMS independent variables

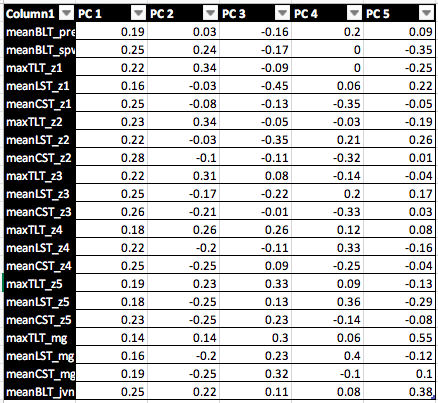

**Table S6**: Table showing the five principal components for Oregon and their loadings in term of ROMS independent variables

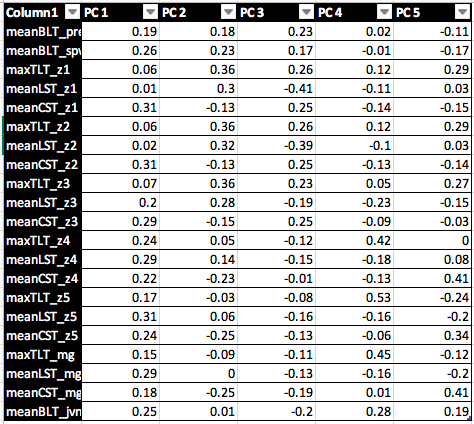

**Table S7**: Table showing the five principal components for Northern California and their loadings in term of ROMS independent variables

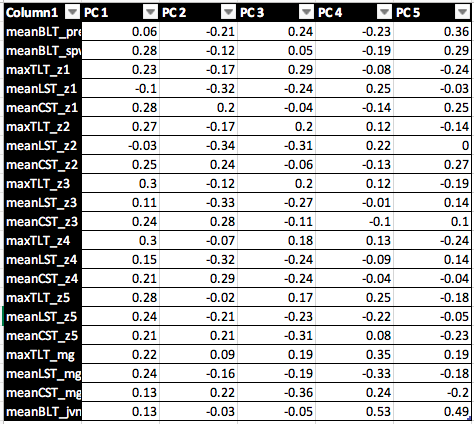

**Table S8**: Table showing the five principal components for Central California and their loadings in term of ROMS independent variables

|  | Washington | Oregon | Northern California | Central California |
| --- | --- | --- | --- | --- |
| PC 1 | 46.85 | 40.69 | 32.7 | 37.66 |
| PC 2 | 22.07 | 17.75 | 23.1 | 26.62 |
| PC 3 | 9.08 | 13.93 | 12.4 | 12.67 |
| PC 4 | 5.96 | 11.13 | 9.38 | 8.02 |
| PC 5 | 4.07 | 4.33 | 6.88 | 4.65 |

**Table S9**: Percentage of variance explained by the five principal components for each region.

| **Model** | **Intercept** | **PC1** | **PC2** | **PC3** | **R^2^**  **whole**  **data set** | **AICc**  **whole**  **data set** | **ΔAICc**  **whole**  **data set** |
| --- | --- | --- | --- | --- | --- | --- | --- |
| 1 | 2.045 |  |  |  | 0.000 | 35.9 | 0.00 |
| 2 | 2.045 |  |  | 0.078 | 0.070 | 36.2 | 0.30 |
| 3 | 2.045 | 0.031 |  |  | 0.058 | 36.6 | 0.66 |
| 4 | 2.045 | 0.031 |  | 0.078 | 0.128 | 36.9 | 1.02 |
| 5 | 2.045 |  | -0.026 |  | 0.019 | 37.8 | 1.89 |

**Table S10**: Results of the Washington model selection using the five principal components on whole data set showing models with $\Delta AICc<2$.

| **Model** | **Intercept** | **PC1** | **PC2** | **PC4** | **R^2^**  **whole**  **data set** | **AICc**  **whole**  **data set** | **ΔAICc**  **whole**  **data set** |
| --- | --- | --- | --- | --- | --- | --- | --- |
| 1 | 1.915 | -0.075 |  | -0.146 | 0.436 | 32.8 | 0.00 |
| 2 | 1.915 | -0.075 | -0.040 | -0.146 | 0.463 | 34.2 | 1.44 |

**Table S11**: Results of the Oregon model selection using the five principal components on whole data set showing models with $\Delta AICc<2$.

| **Model** | **Intercept** | **PC1** | **PC2** | **PC4** | **R^2^**  **whole**  **data set** | **AICc**  **whole**  **data set** | **ΔAICc**  **whole**  **data set** |
| --- | --- | --- | --- | --- | --- | --- | --- |
| 1 | 1.343 | -0.0781 | -0.096 | -0.161 | 0.4048 | 49.8 | 0.00 |

**Table S12**: Results of the Northern California model selection using the five principal components on whole data set showing models with $\Delta AICc<2$.

| **Model** | **Intercept** | **PC1** | **PC4** | **PC5** | **R^2^**  **whole**  **data set** | **AICc**  **whole**  **data set** | **ΔAICc**  **whole**  **data set** |
| --- | --- | --- | --- | --- | --- | --- | --- |
| 1 | 0.403 | -0.132 | -0.222 | 0.444 | 0.5518 | 65 | 0.00 |

**Table S13**: Results of the Central California model selection using the five principal components on whole data set showing models with $\Delta AICc<2$.

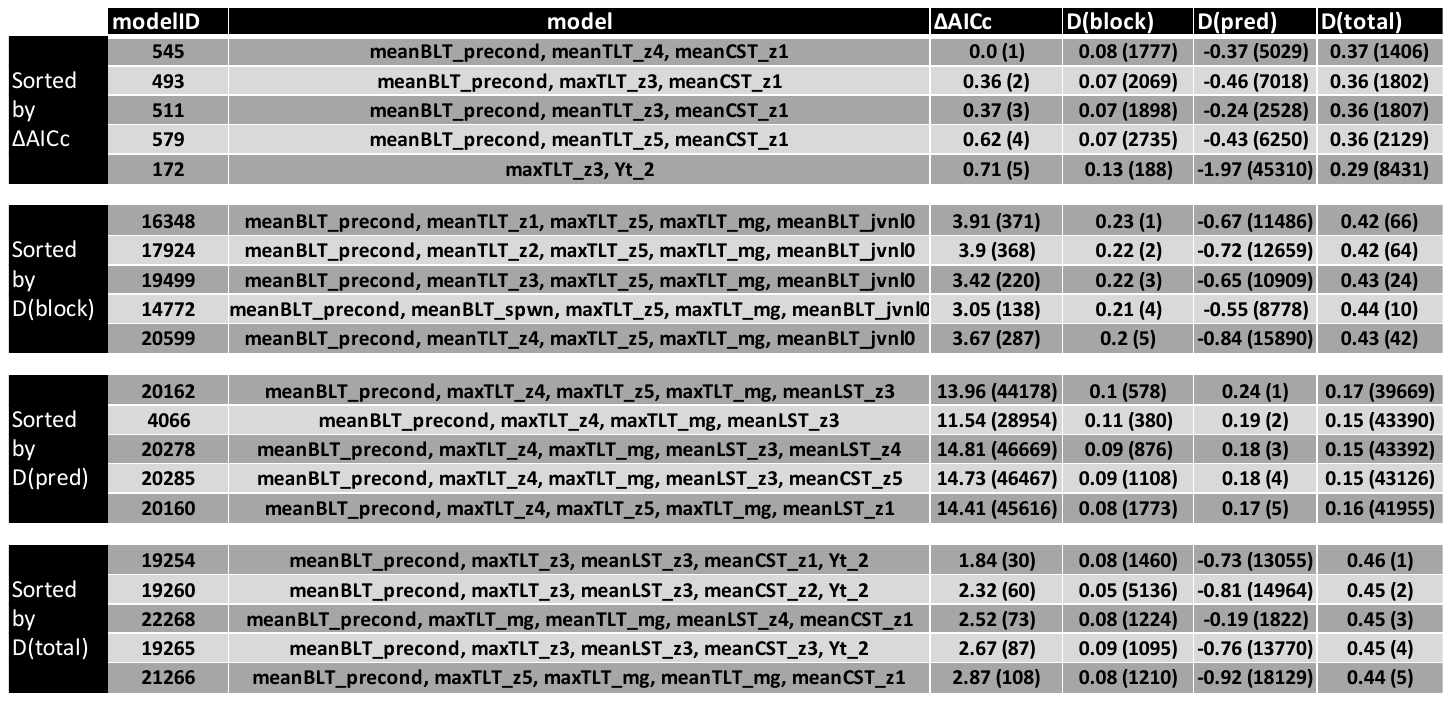

**Table S14:** Top five models for each of the four metrics for Washington for the models with lagged variables. The results show the value of the metric and its rank (between parentheses). The ΔAICc is ranked increasing order and D(pred), D(block) and D(total) are ranked in decreasing order.

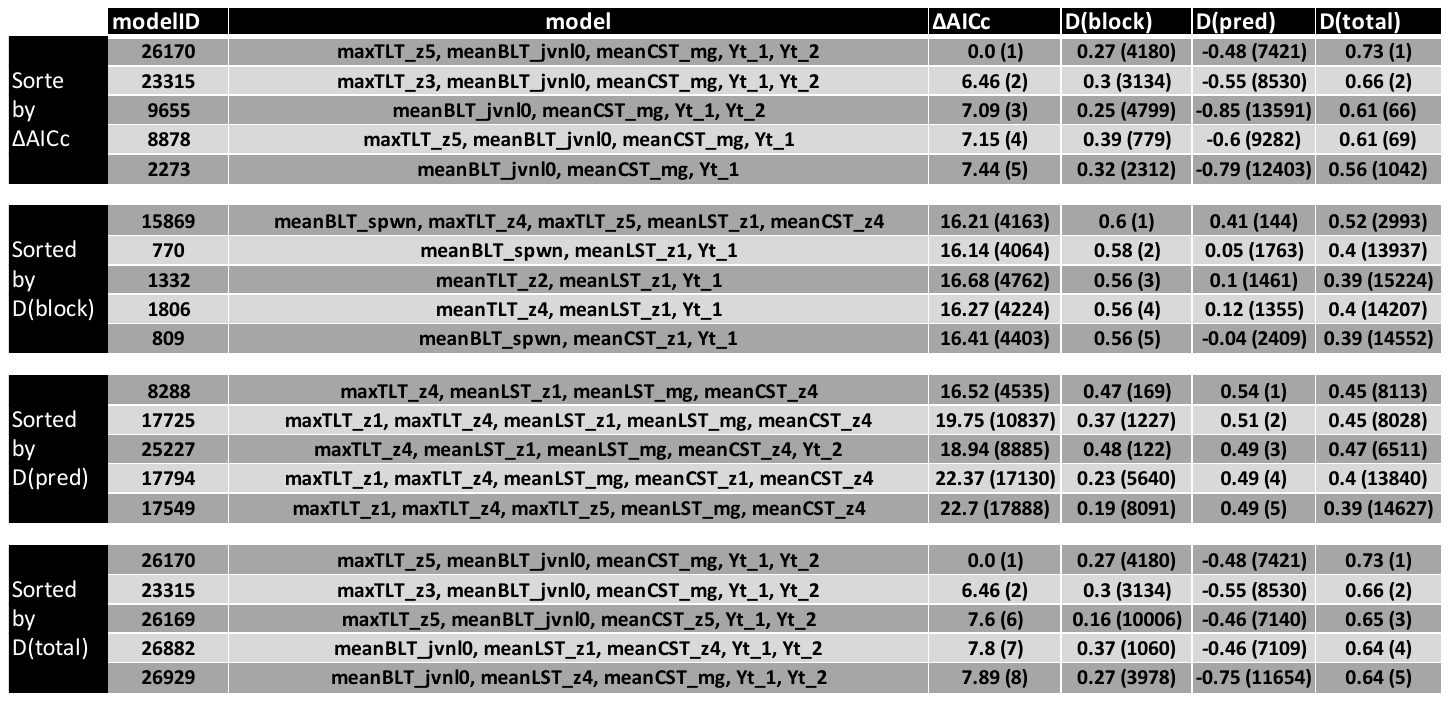

**Table S15:** Top five models for each of the four metrics for Oregon for the models with lagged variables. The results show the value of the metric and its rank (between parentheses). The ΔAICc is ranked increasing order and D(pred), D(block) and D(total) are ranked in decreasing order.

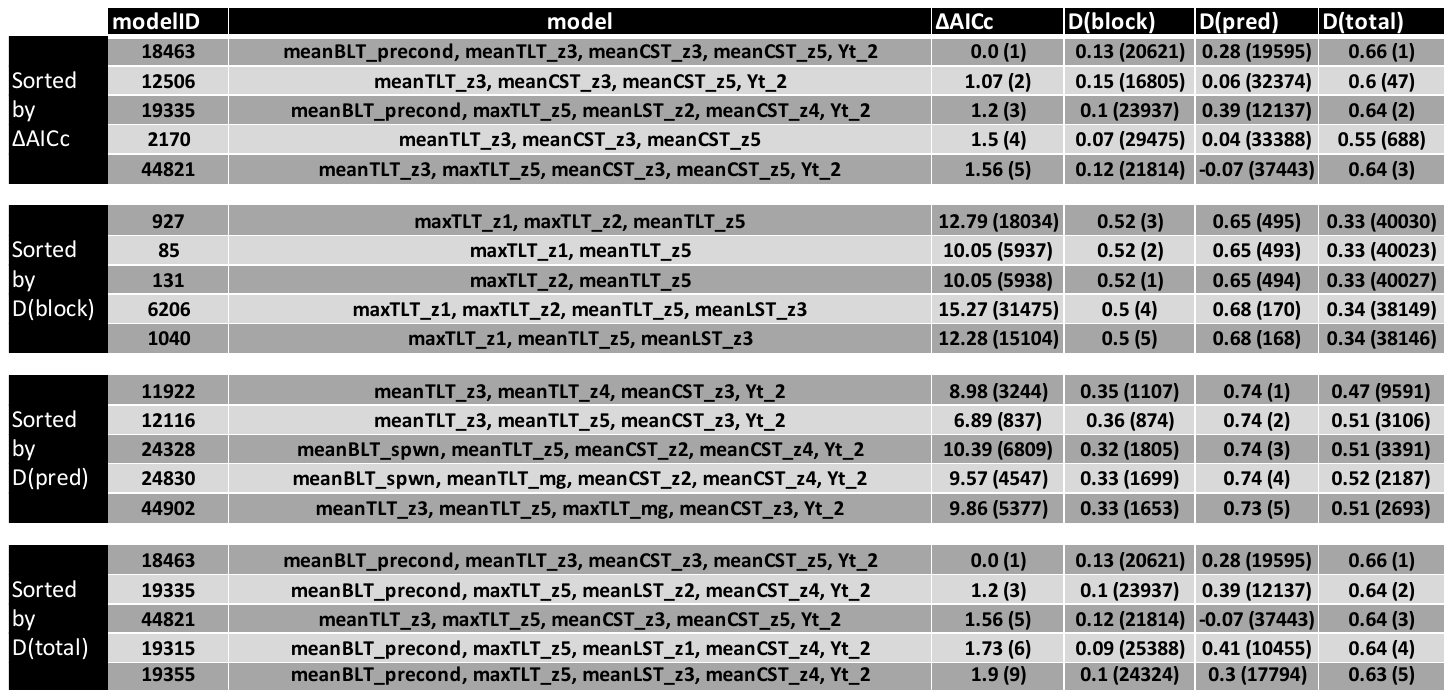

**Table S16:** Top five models for each of the four metrics for Northern California for the models with lagged variables. The results show the value of the metric and its rank (between parentheses). The ΔAICc is ranked increasing order and D(pred), D(block) and D(total) are ranked in decreasing order.

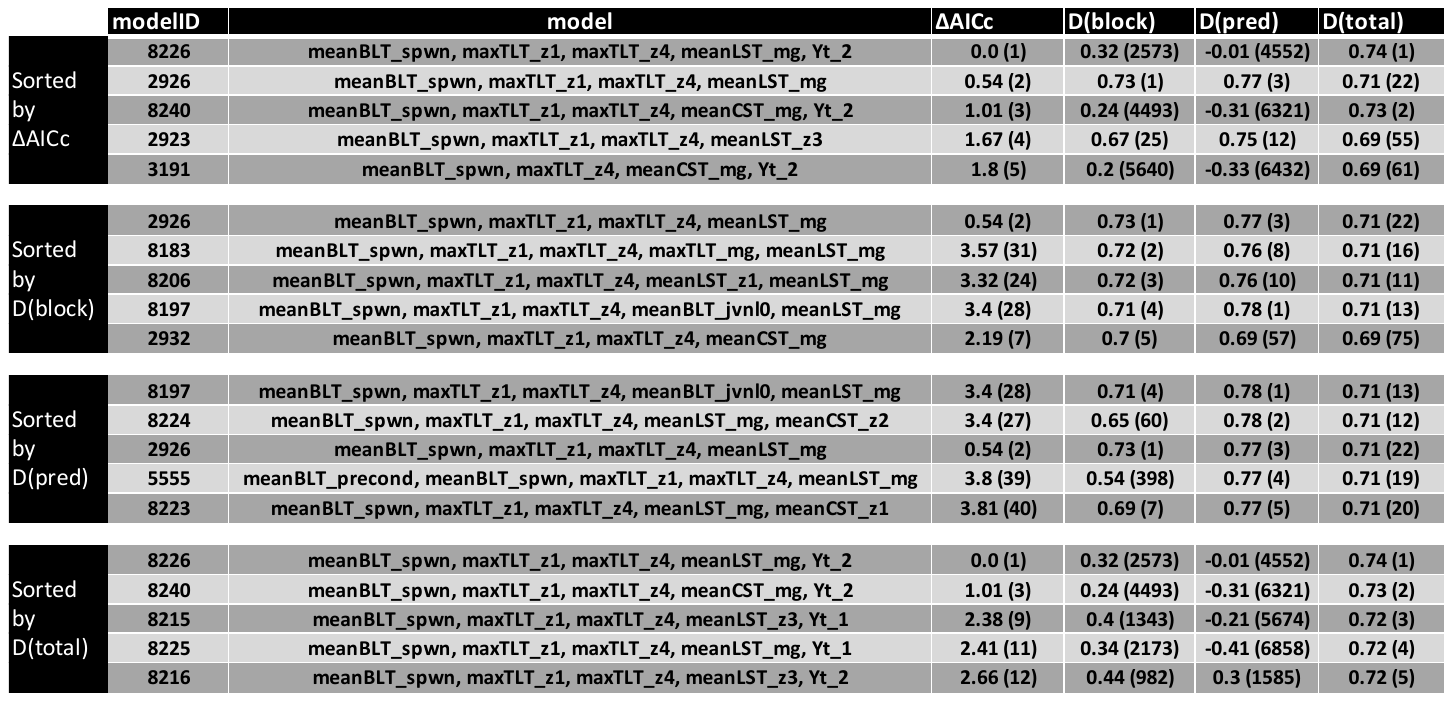

**Table S17:** Top five models for each of the four metrics for Central California for the models with lagged variables. The results show the value of the metric and its rank (between parentheses). The ΔAICc is ranked increasing order and D(pred), D(block) and D(total) are ranked in decreasing order.

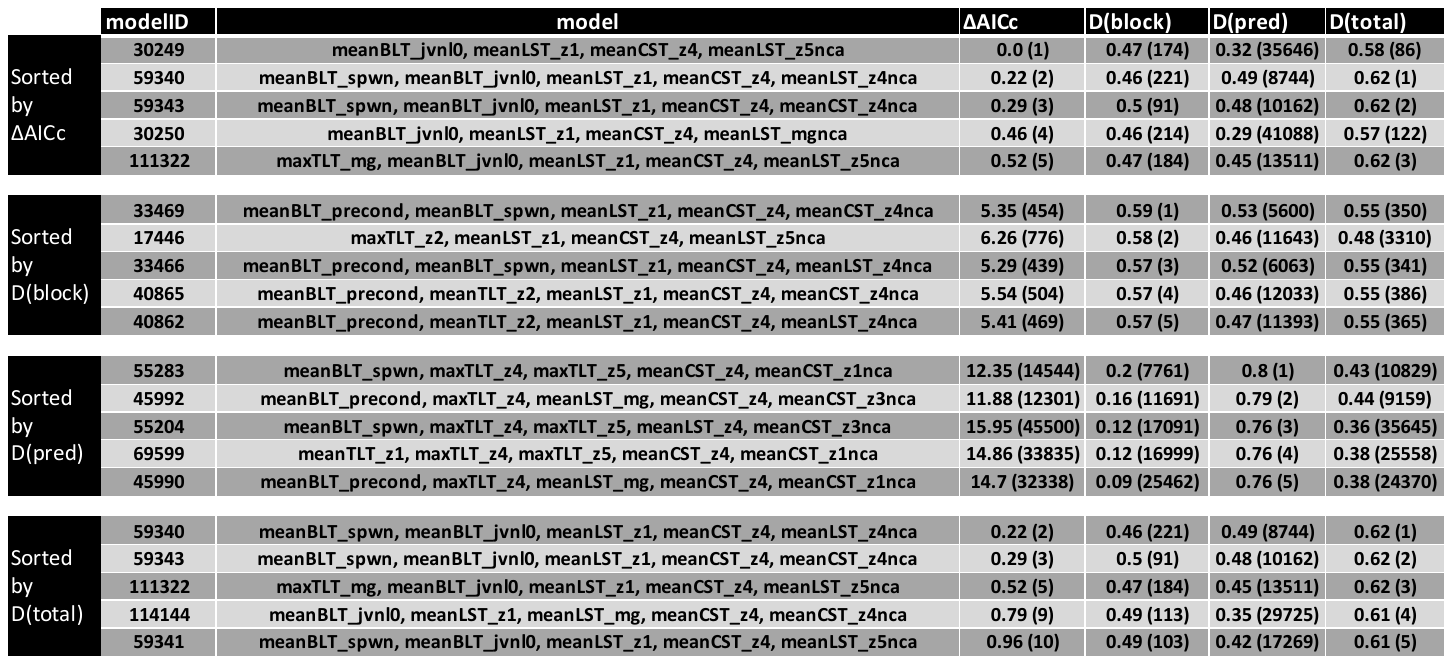

**Table S18:** Results of model selection showing the top five models for each of the four model selection metrics when transport variables from Northern California are considered as predictors of preseason abundance for Oregon. The results show the value of the metric and its rank (between parentheses). The ΔAICc is ranked increasing order and D(pred), D(block) and D(total) are ranked in decreasing order.

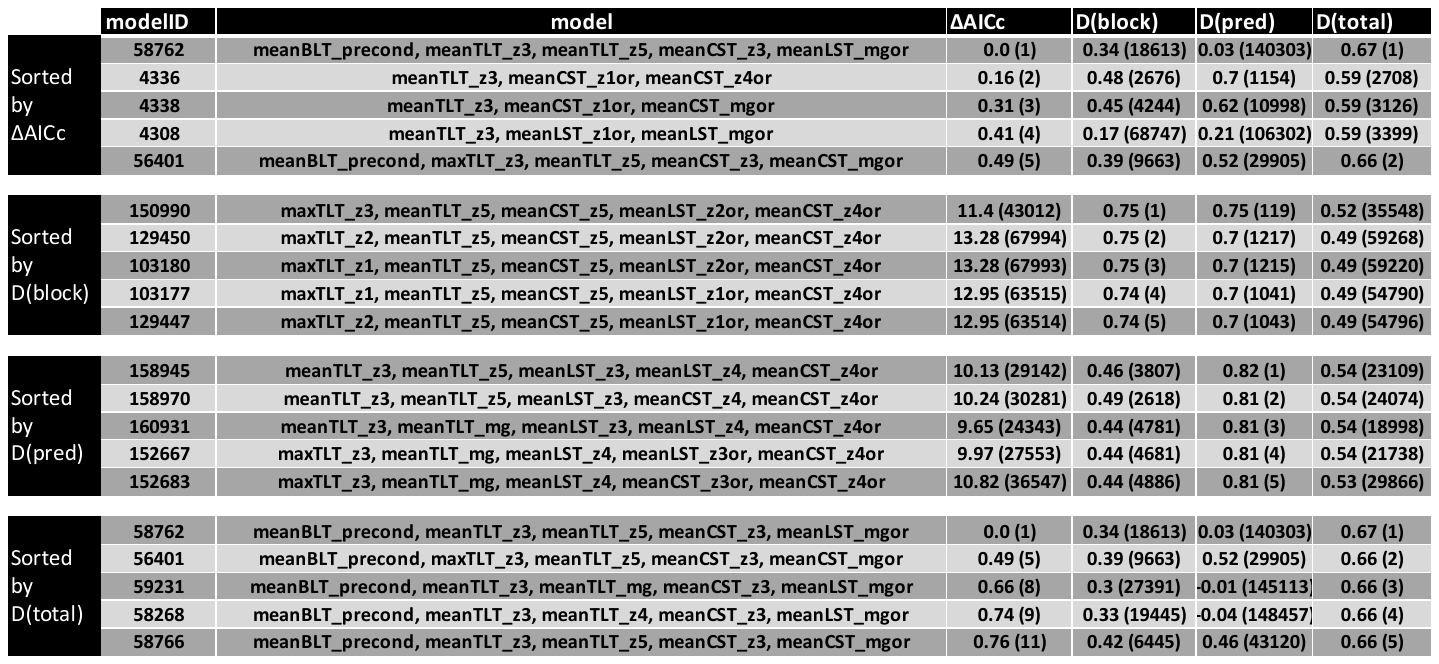

**Table S19:** Results of model selection showing the top five models for each of the four model selection metrics when transport variables from Oregon are considered as predictors of preseason abundance for Northern California. The results show the value of the metric and its rank (between parentheses). The ΔAICc is ranked increasing order and D(pred), D(block) and D(total) are ranked in decreasing order.

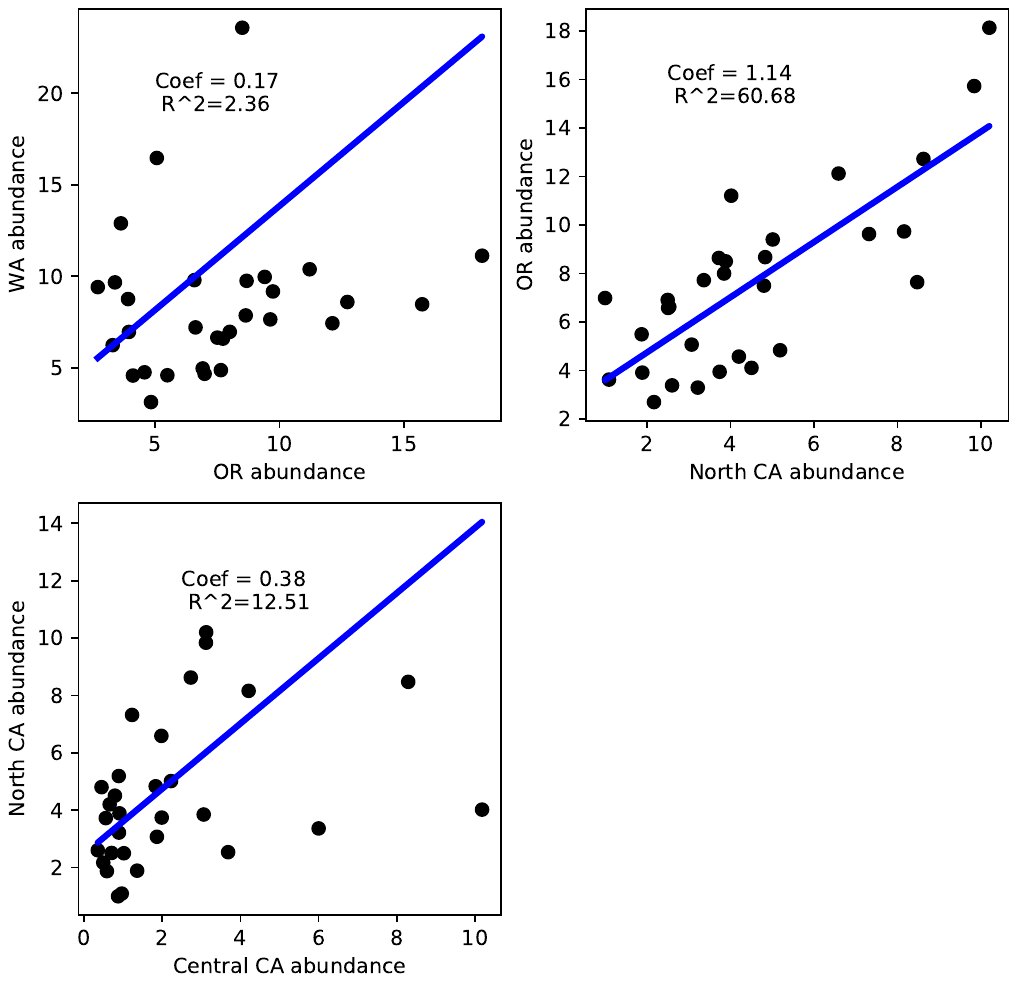

**Figure S10**: Correlations between legal-sized male Dungeness crab abundances from the different regions.

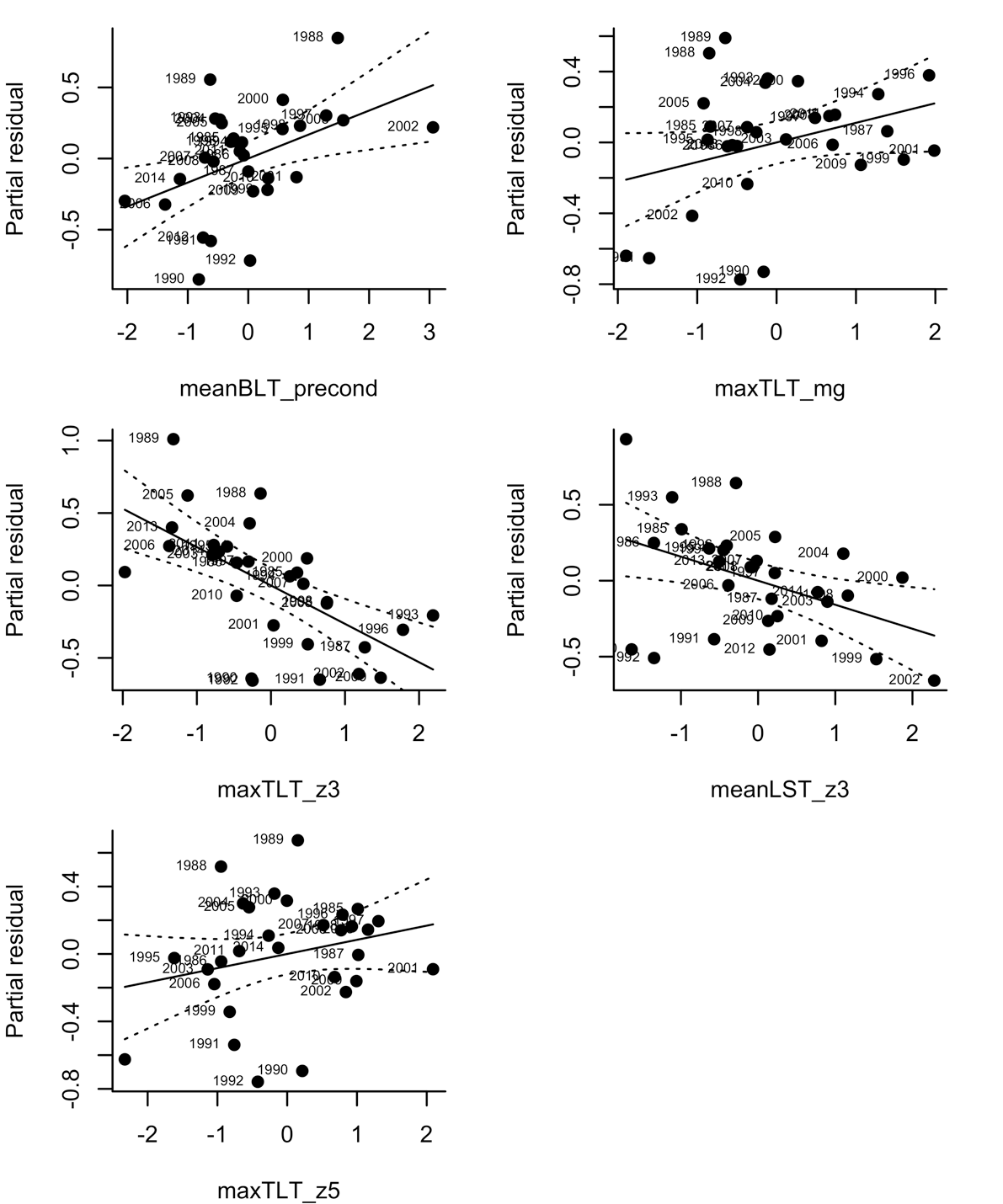

**Figure S11**: Partial residual plots for the independent variables in the best-fit model for Washington.

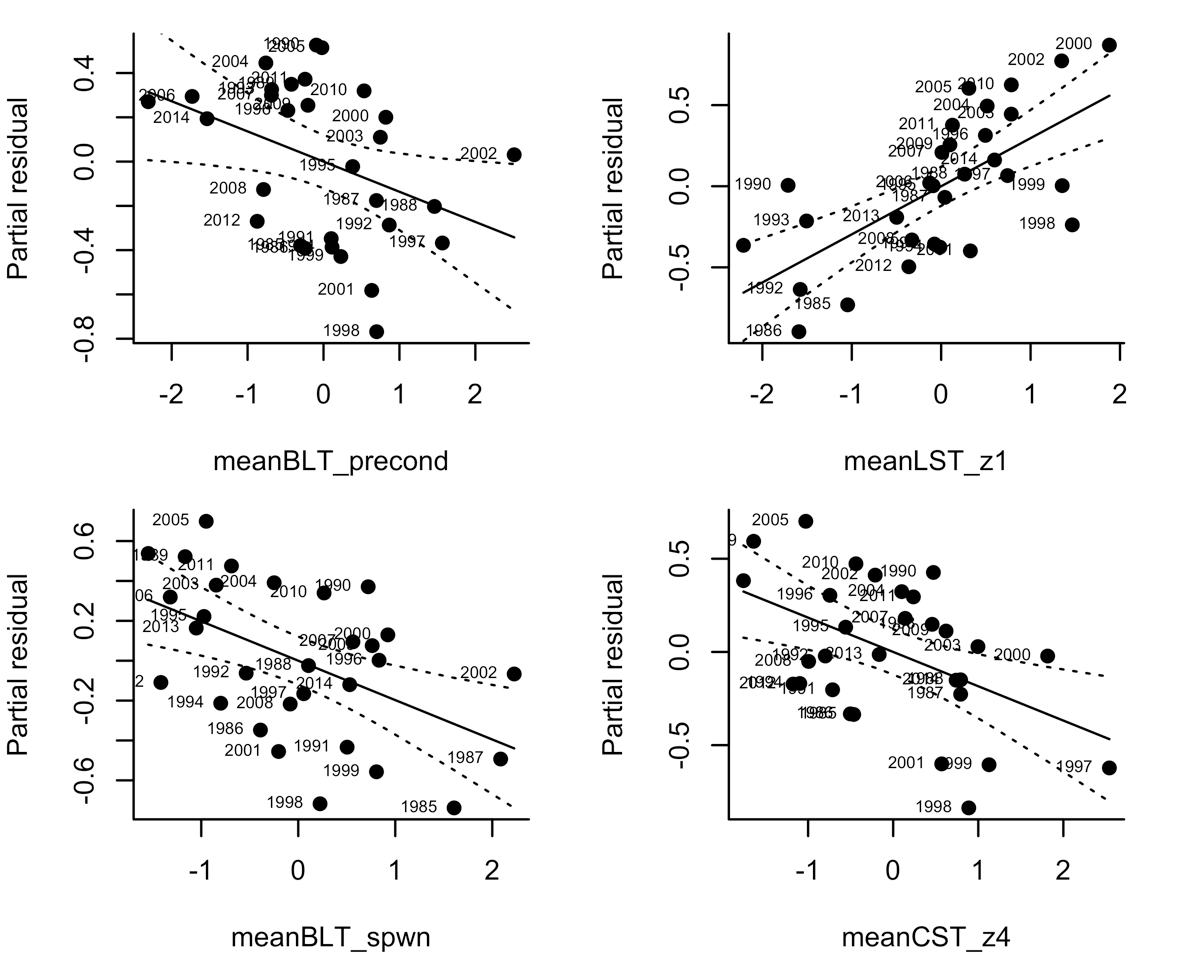
**Figure S12**: Partial residual plots for the independent variables in the best-fit model for Oregon.

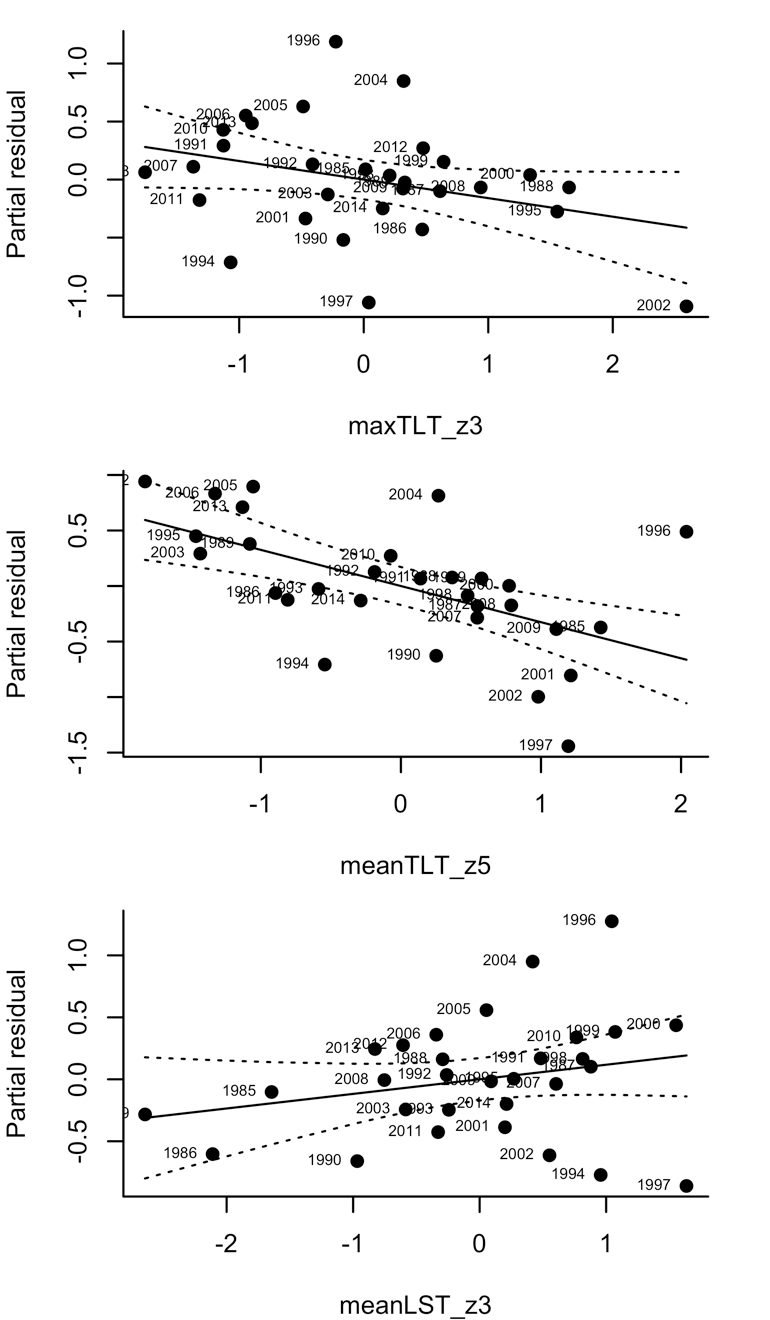
**Figure S13**: Partial residual plots for the independent variables in the best-fit model for Northern California.

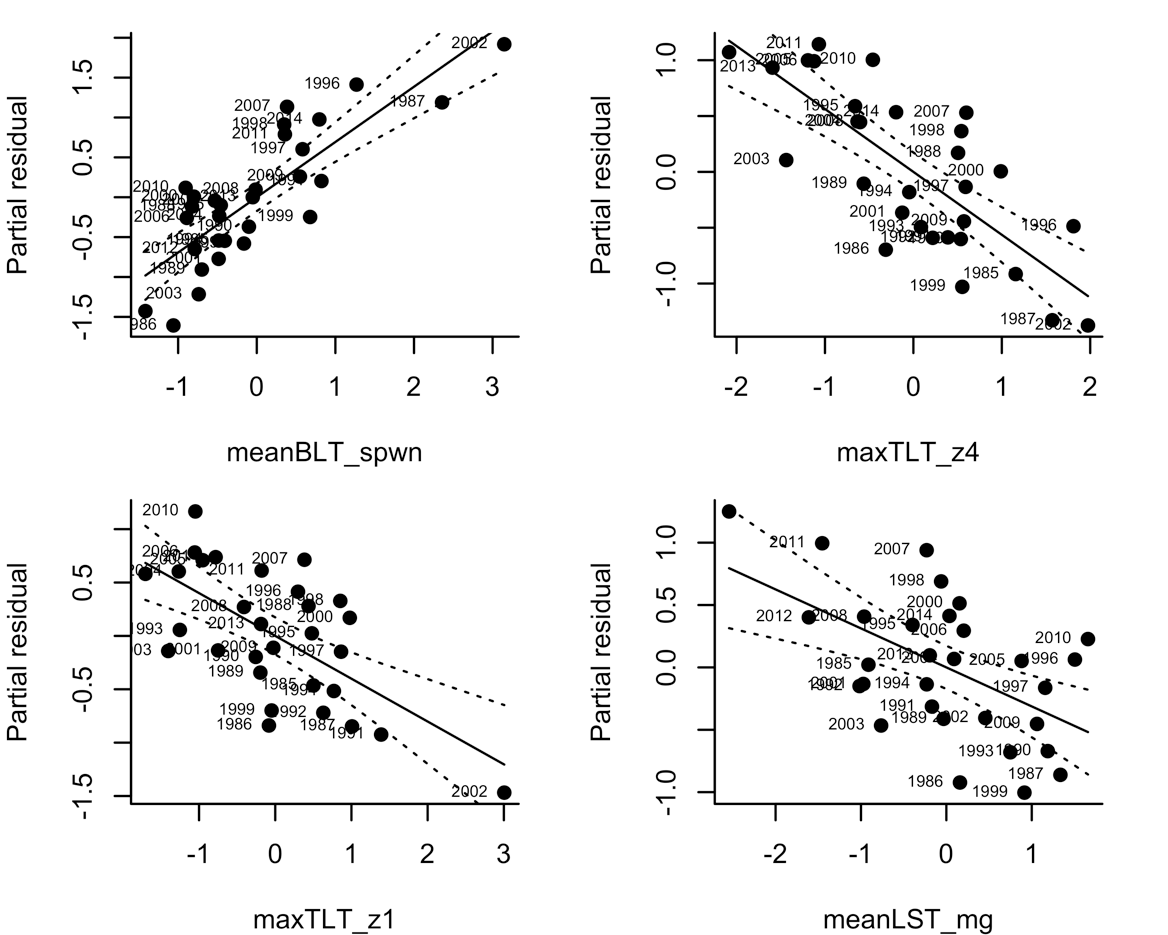
**Figure S14**: Partial residual plots for the independent variables in the best-fit model for Central California.

**Figure S15**: (a) frequency distribution of R^2^ values from the jackknife resampling for individual years. (b) R^2^ when the indicated year was removed from fitting of the best-fit model for Washington

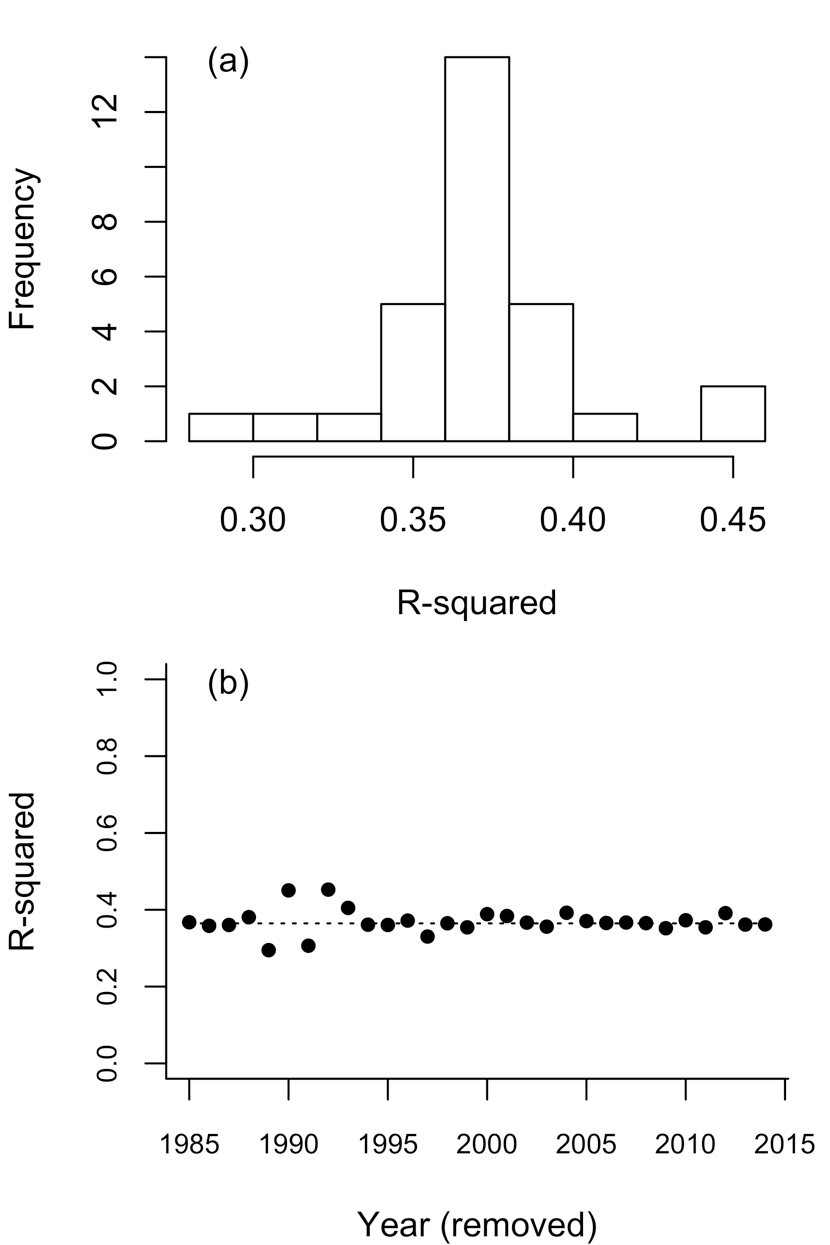

**Figure S16**: (a) frequency distribution of R^2^ values from the jackknife resampling for individual years. (b) R^2^when the indicated year was removed from fitting of the best-fit model for Oregon

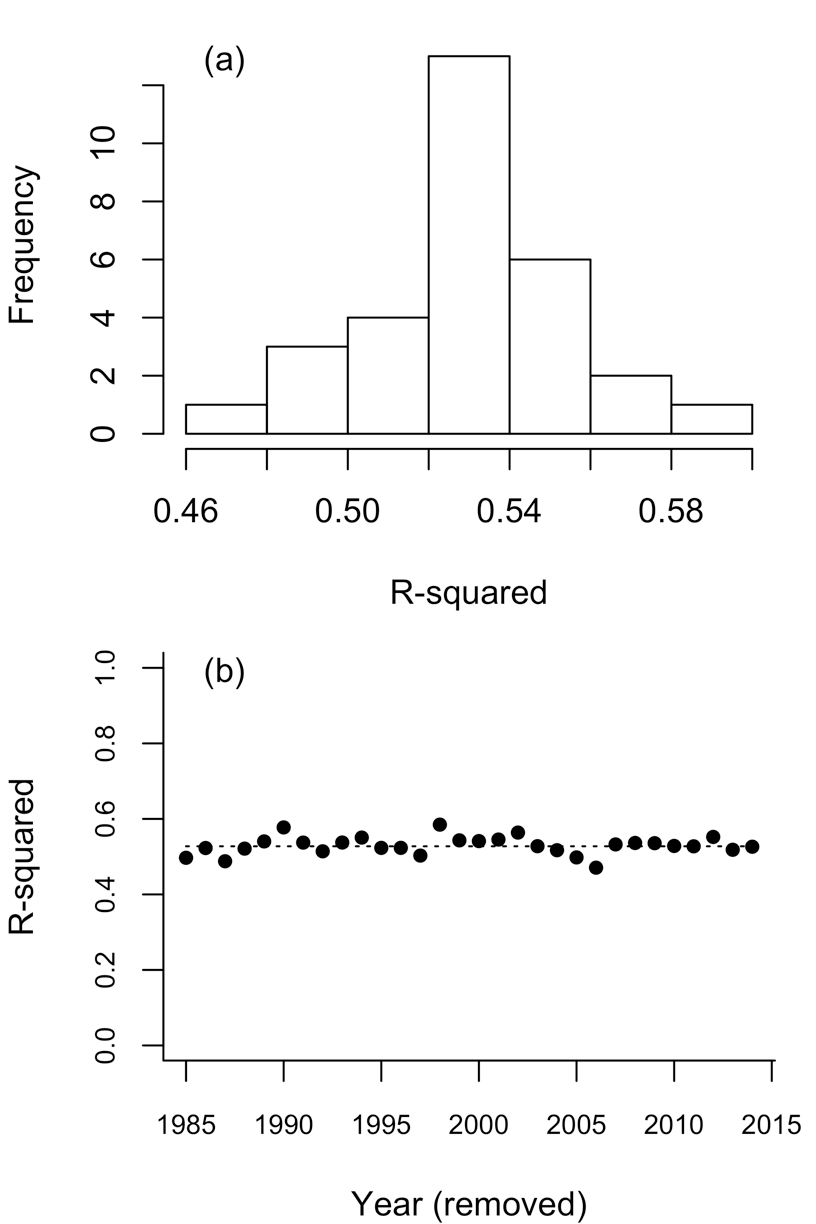

**Figure S17**: (a) frequency distribution of R^2^ values from the jackknife resampling for individual years. (b) R^2^ when the indicated year was removed from fitting of the best-fit model for Northern California.

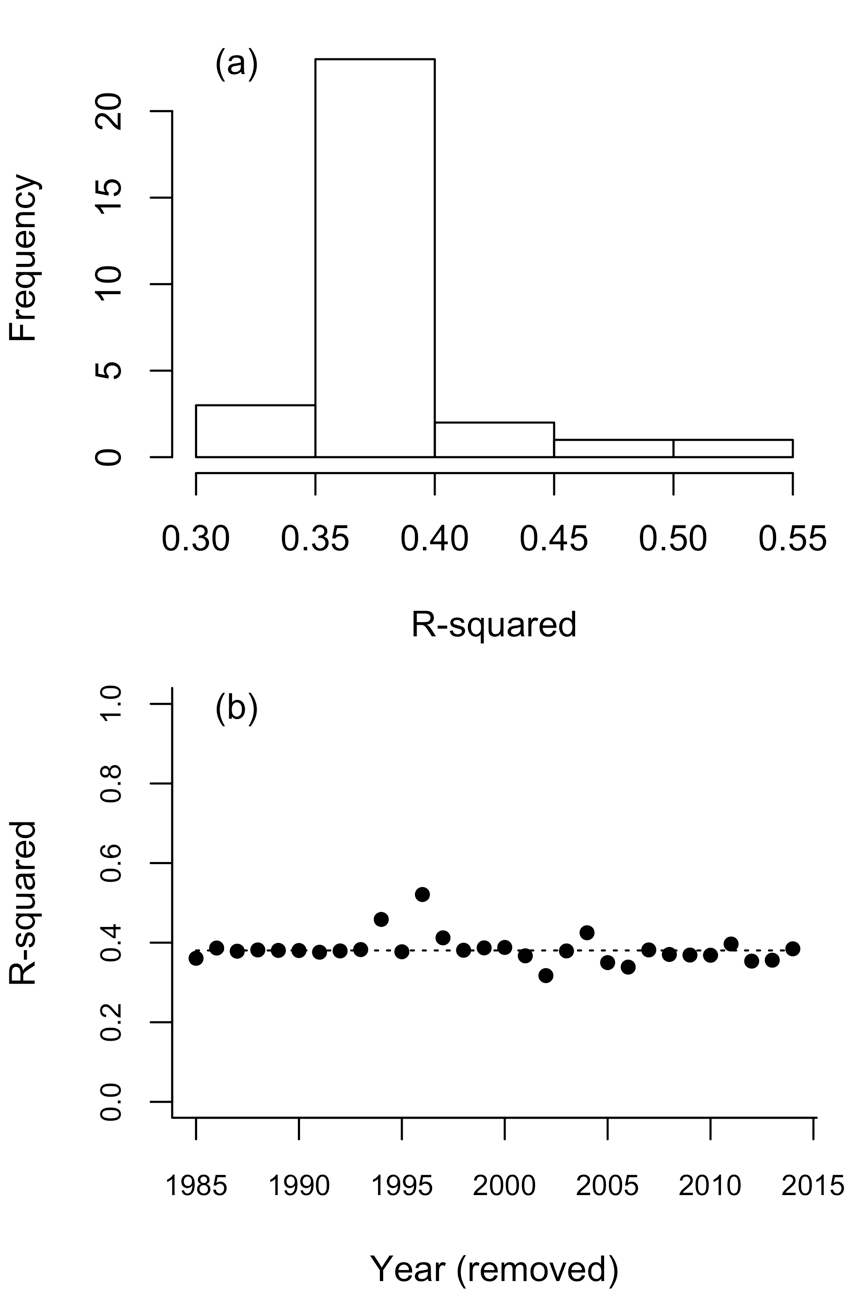

**Figure S18**: (a) frequency distribution of R^2^ values from the jackknife resampling for individual years. (b) R^2^ when the indicated year was removed from fitting of the best-fit model for Central California.

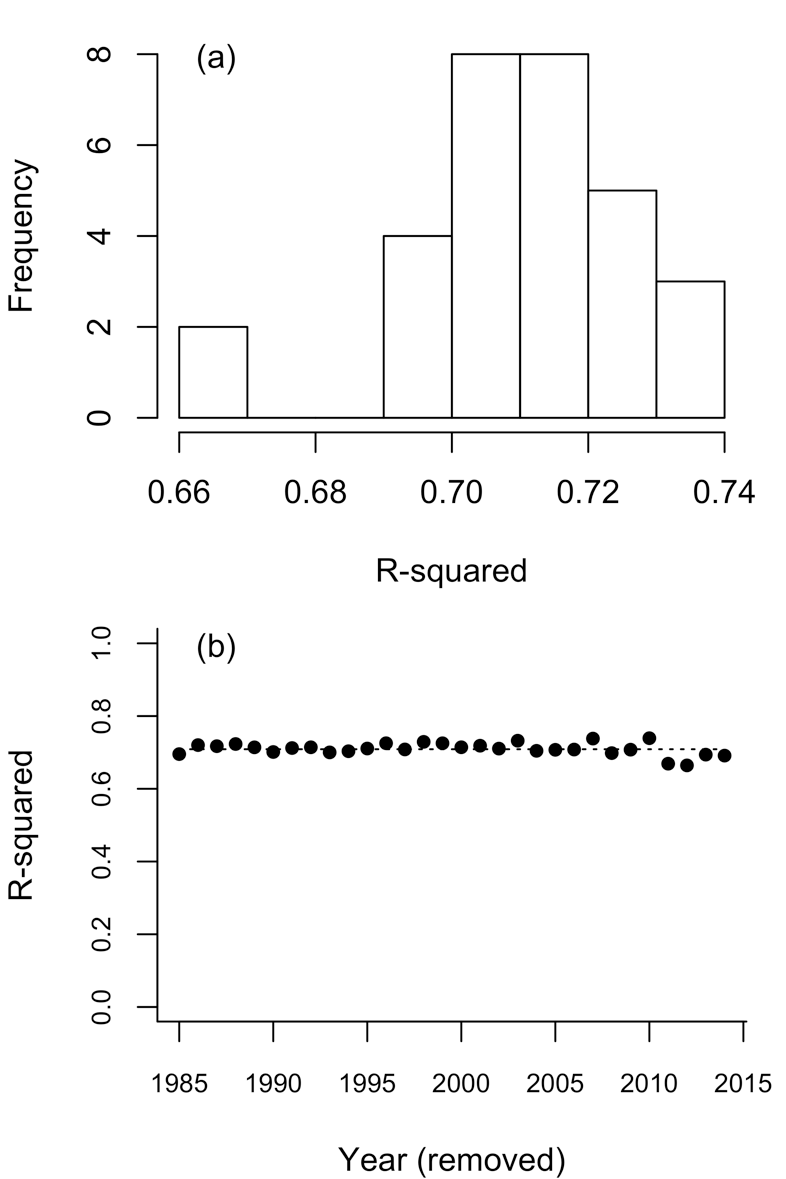
